# Optimization and redevelopment of single-cell data analysis workflow based on deep generative models

**DOI:** 10.1101/2022.09.12.507562

**Authors:** Yunhe Liu, Qiqing Fu, Chenyu Dong, Xiaoqiong Xia, Gang Liu, Lei Liu

## Abstract

The present single-cell RNA sequencing(scRNA-seq) analysis pipelines require a combination of appropriate normalization, dimension reduction, clustering, and specific-gene analysis algorithms, but the rationale for the choice of these algorithms is relatively subjective because of the lack of ground truth assessment conclusions. As the number of captured single-cells increases, the number of different types of noise cells also increases, which can strongly affect the analysis efficiency. For scRNA-seq, a technology that generates data through multi-process operations, the deep generative model should be a good choice for this type of data analysis, allowing simultaneous estimation of multiple unobservable parameters assumed in the data generation process. Hence, in our study, we sequenced a pool of pre-labeled single cells to obtain a batch of scRNA-seq data with main and fine labels, which was then used to evaluate the clustering and specific-gene analysis methods. Afterward, we applied two deep generative models to infer the probabilities of pseudo and impurity cells. And by stepwise removing the inferred noise cells, the clustering performance and the consistency of different specific-gene analysis methods are both greatly improved. After that, we applied Deep-LDA (a latent Dirichlet allocation-based deep generative model) to scRNA-seq data analysis. And this model takes the count matrix as input, and makes the classification and specific gene optimization process mutually dependent, which has more practical sense and simplifies the analysis workflow. At last, we successfully implemented the model with transferred knowledge to make single-cell annotation and verified its superior performance.

## Introduction

ScRNA-seq has grown to be the primary approach for research on histo-differentiation^1^, brain structure^2^, and tumor immunology^3, 4, 5, 6^. However, in practice, the outcomes generated from this data type are frequently unstable^7^. The major reason is that the obtained data which was sequenced from a huge number of single cells, is insufficient to cover the whole transcriptome feature of each cell, leading to a high 0-inflation ratio and technical noise^8^. Moreover, the transcripts of different cell types have poor overlap, causing a more complicated distribution in observed data.

Many algorithms have been developed to reduce the noise within and between the cells. BASiCS^9^ and Scran^10^ estimate a specific scale factor for normalization. MNN algorithm^11^ has been employed to integrate data from multiple batches. DCA^12^, scVI^13^, and ZINB-WaVE^14^ infer noise-free expression values with generative models. And SCTransform^15^ corrects the dependence between gene expression values and sequencing depths by means of regularized negative binomial distribution regression. The gains from these methods are remarkable for the current clustering and specific gene analysis. But for these methods reshaping observed expression value, there is a high possibility of eliminating some useful information when increasing the positive signal in their assumption.

The commonly used single-cell clustering procedures work on the cell nearest neighbor network by KNN and SNN, and then utilize the community-detection algorithms^16^, such as louvain^17^ and leiden^18^, to iteratively optimize cell clusters. And the existed noise cells which distributed widely separated from the target cells in latent space, such as pseudo cells that capture environment RNA^19^ and bits of impurity cells, can largely affect cell neighbor relationship estimation and clustering iteration. Thus, removing these noise cells will theoretically improve the performance of the current clustering algorithms. The specific gene analysis depends on clustering analysis, which in turn is not. That is to say, the quality of clustering will directly affect the specific genes’ quality, while the generated class-specific genes cannot reversely optimize the clustering. However, in the practical sense, the cell-specific genes should be the law for their sorting. So, it will be more meaningful to enable the two analysis steps to be coupled and mutually dependent.

The scRNA-seq data analysis pipelines consist of several steps^20^, each of which has many optional algorithms to choose from^21^. Due to lake of suitable ground truth, it’s hard to objectively evaluate these pipelines^16, 22^. Although the simulation of scRNA-seq data drawing from Splatter^8^, SPARSim^23^, SymSim^24^, etc., can produce the pseudo-ground truth for algorithm evaluation, they only reproduce the limited assumed features of the actual data. Instead, the real pre-labeled pooled sample scRNA-seq data will be a more optimal choice of ground truth.

Deep generative models^13^ can combine assumed multiple probability distributions and deterministic relations to generate observed data, and infer the unobservable parameters in the generation process by the variation inference or MCMC^13^. Various types of controlling parameters, such as scale factor, real RNA signal, noise cell probability, etc., can be incorporated into the model generation process and inferred simultaneously. Such models have been shown to be highly suitable for fitting the scRNA-seq data which have complicated features^25, 26, 27^. And some modules of generation are constructed based on the neural network architecture, which have a higher ability to fit the nonlinear elements in data^28, 29^.

Regarding the above issues, we experimentally obtained a batch of scRNA-seq data with main and fine labels and used it as ground truth to evaluate the analysis pipelines. And then, we used deep generative models for the inference of two types of noise cells and used an LDA-based deep generative model^30^ for cell classification and class-specific gene analysis. And finally, we validated the models’ superior performance and developed more related applications.

### A batch of sequencing data from pre-enriched labeled single-cell

Following the procedures of pre-enrichment, labeling, and then pooled sequencing, we obtained a batch of scRNA-seq data from mixed 10 types of immune cells and a batch of bulk-seq data from 8 of them (Fig 1a; Methods). After preprocessing, dimension reduction and visualization of the two types of data (Methods), the distribution of the cell types in scRNA data (Fig 1b) is spatially consistent with that in bulk-seq data (Fig 1d left). Subsequently, we employed CIBERSORT^31^ to decompose the bulk-seq data (Fig.1d right, Methods), and also verified the consistency. After that, we collected 460 specific genes (Methods; Table S8) from 3 databases and 4 references for the 10 types of immune cells. And the differential analysis verified that most of the collected genes were significant in both types of sequencing data, except for some sub-categories of T cells (Fig 1c; Table S8). The heatmap of robust specific genes in the two types of data (Fig 1b; Fig S1a) and the UMAP of some key gene intensities (Fig S1b) showed that the genes have high specificity in corresponding cell types. In summary, we suggested that the batch of scRNA-seq data with main and fine labels can be used as ground truth for classification algorithm evaluation.

**Figure 1.**
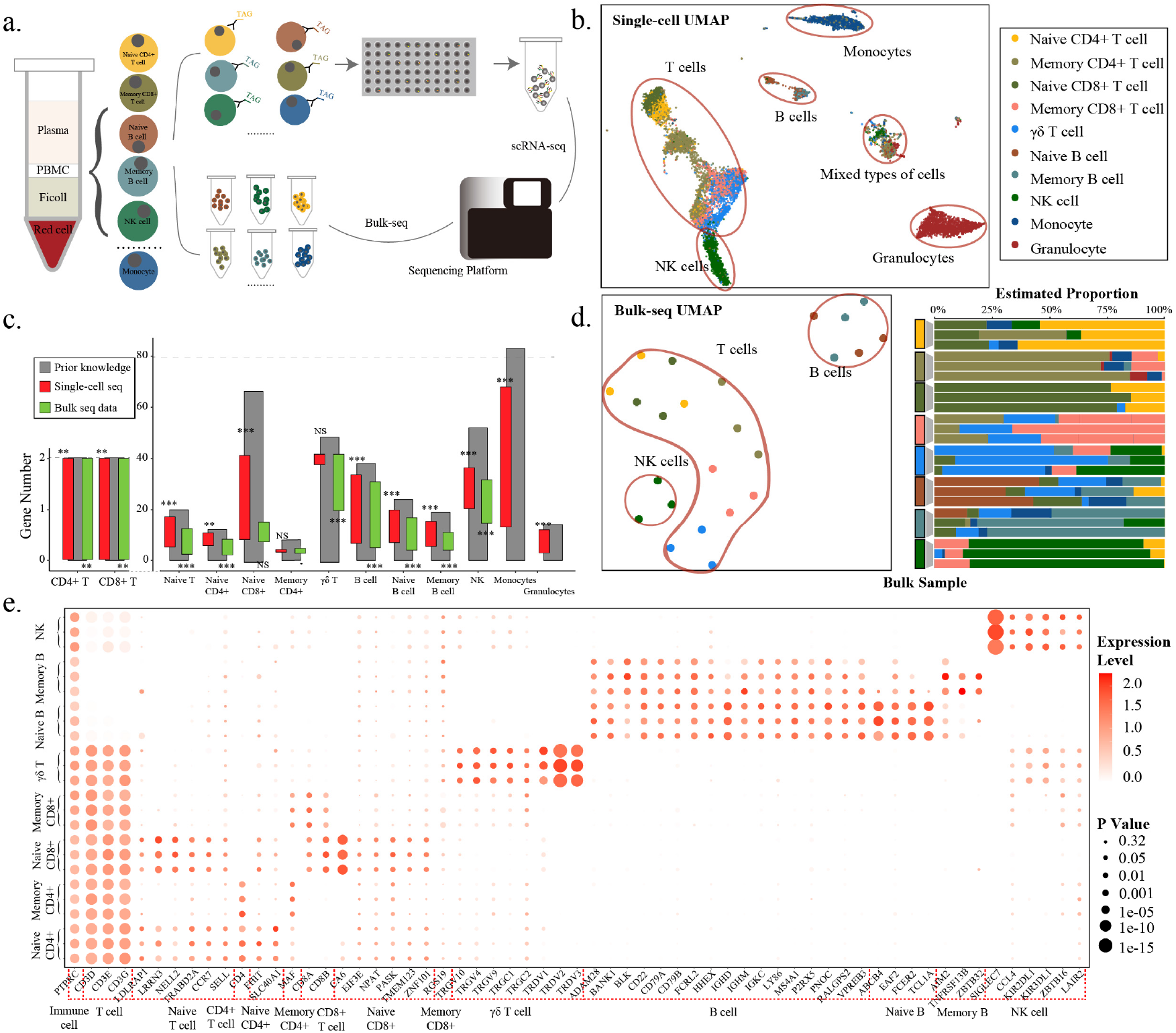
Analysis of pre-labeled immune cell sequencing data **a**. Schematic diagram of the sequencing process of pre-enriched labeled immune cells; **b**. UMAP visualization of the pooled scRNA-seq data, color-coded labels for pre-enriched cell types; **c**. Bar plot of the specific genes in prior knowledge (grey) and two small bars showing which fraction is significant after differential analysis (**Table S8**) in single-cell data (red) and bulk data (green), respectively (the horizontal overlap between the three bars is the intersection domain). Independence tests (fisher test) were applied to examine the relationship between the differential analysis results and the prior knowledge, and its significance was marked above or below the corresponding bar (‘***’, ‘**’, ‘*’, ‘·’, ‘NS’ represent P values <0.001, <0.01, <0.05, <0.1, >0.1, respectively); **d**. UMAP visualization of bulk sequencing data (left panel) and estimated cell-type component proportion (right panel). The color panel for cell types is the same as **a**; **e**. Robust specific-gene heatmap for bulk sequencing data.

### Clustering methods comparison and improvement after inferred noise cell removal

After removing the inferred pseudo-cells and impurity cells by two deep generative models, the remaining data were visualized by UMAP^32^ and tSNE^33^ (Fig 2a; Fig S2a, b; Methods). The cell spatial distribution showed that the mixed-type cell distributed area (Fig 1b) was inferred as pseudo-cells (Fig 2a left; Fig S2b left), which were composed of the same ambient RNA under different cell type labels, and some small patches of cells sporadically distributed in the plot (Fig 2a middle; Fig S2b middle) were inferred as impurity cells. And with stepwise removal of the noise cells, the aggregation degrees for different types of cells were enhanced, especially for Memory CD8+ T cells (Fig 2a). Then, we applied the 3-phase data to evaluate the effectiveness of 5 single-cell clustering procedures with 13 clustering metrics (Methods). And the evaluation results indicated that the noise cell removal significantly improved the clustering scores for all procedures (Fig 2b; Fig S3a-f). And the overall scores of louvain-, leiden- and densitypeak^34^-based clustering were outstanding and comparable, which we then suggested as the first echelon methods. While DDRTree^35^- and Consensus^36^-based clustering were seriously affected by noise and the overall scores were lower. In addition, with noise cell removal, the consistency between the distribution of annotated cluster and corresponding real cell type was improved on main cell labels (Fig 2c left, Fig S4), while poorly improved on fine cell labels which present more irregular shape within the main label distributed area, such as Naive CD4+ and Memory CD8+ T cells (Fig 2c right, Fig S4). We deduced that the traditional clustering algorithm based on Euclidean distance is difficult to fit irregularly distributed cells accurately.

**Figure 2.**
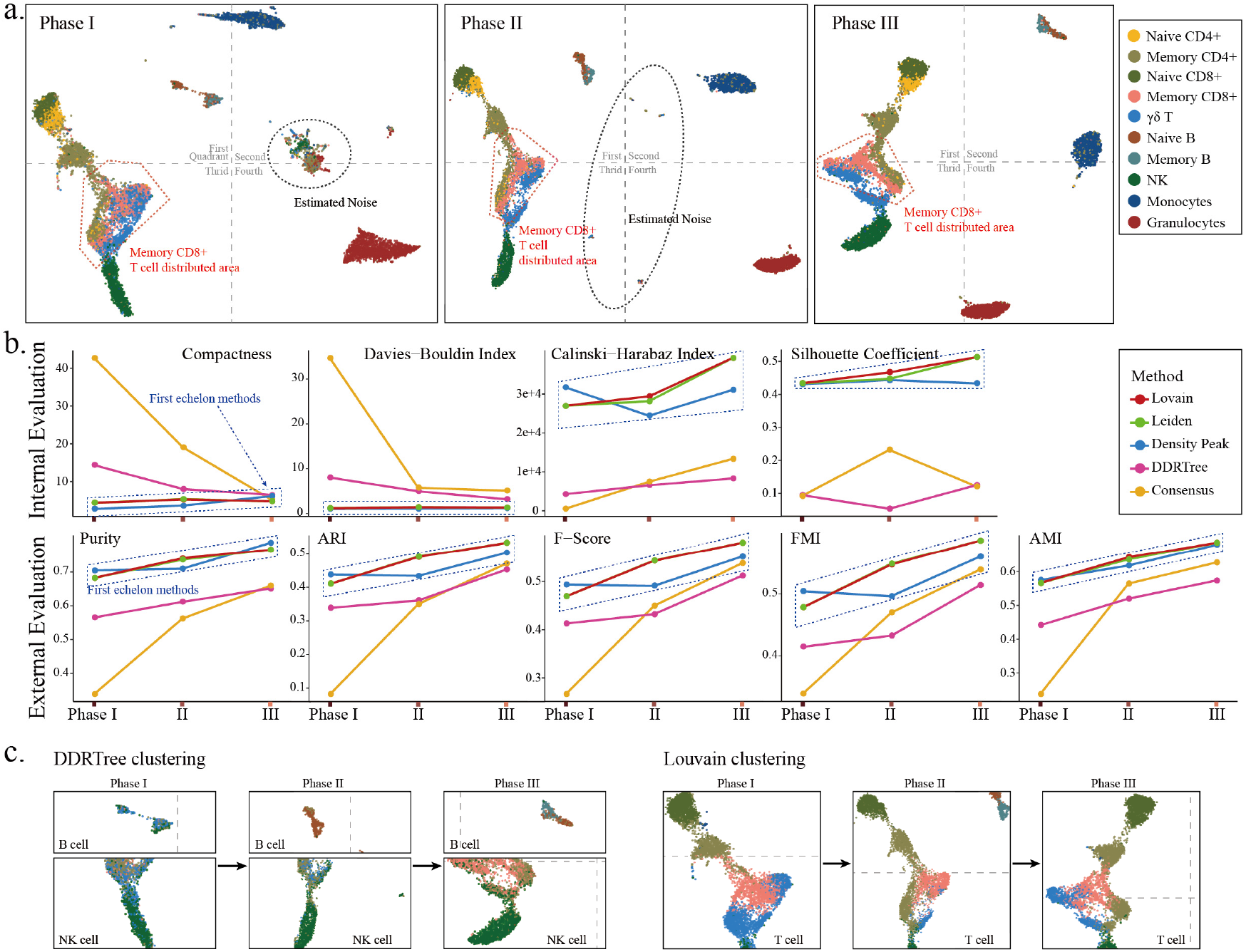
Evaluation of traditional clustering effectiveness after de-noising **a**. UMAP visualization of cells without noise removal (Phase I; 13710 cells left), after removal of inferred pseudo-cells (Phase II; 11319 cells left), after removal of inferred impurity cells (Phase III; 10374 cells left), color-coded labels for pre-enriched cell types; **b**. The line plot of 4 class internal and 5 class external metrics for 5 clustering procedures on the 3-phase data; **c**. Part of the cluster distribution changed following noise cells stepwise removal, and clusters are labeled by the color of the largest proportion of cells and the color panel for cell types is the same as **a**.

### The consistency between different specific-gene analysis procedures

We employed 5 specific-gene analysis procedures to reanalyze the specific genes for the 10 types of immune cells (Methods). The intersection of obtained specific-gene lists showed that t.test-, limma^37^-, and wilcox-based procedures had high consistency (Fig 3a part 2-3), while FC.rank (Fold Change) and LRT-based (Likelihood Ratio Test) procedures had low consistency with all other procedures (Fig 3a part 4-6). And the gene lists from the 3 high consistency methods showed more gene intersection with the prior collected specific-gene lists (Fig 3a part 1; Fig S5; Table S8), which we then nominated as primary methods. Moreover, with the noise cell removal, the overall consistency of the 5 methods was improved (Fig. 3b), which was mainly demonstrated by significantly reduced non-intersection part (Fig. 3b left 1) and significantly increased co-intersection part (Fig 3b right 1).

**Figure 3.**
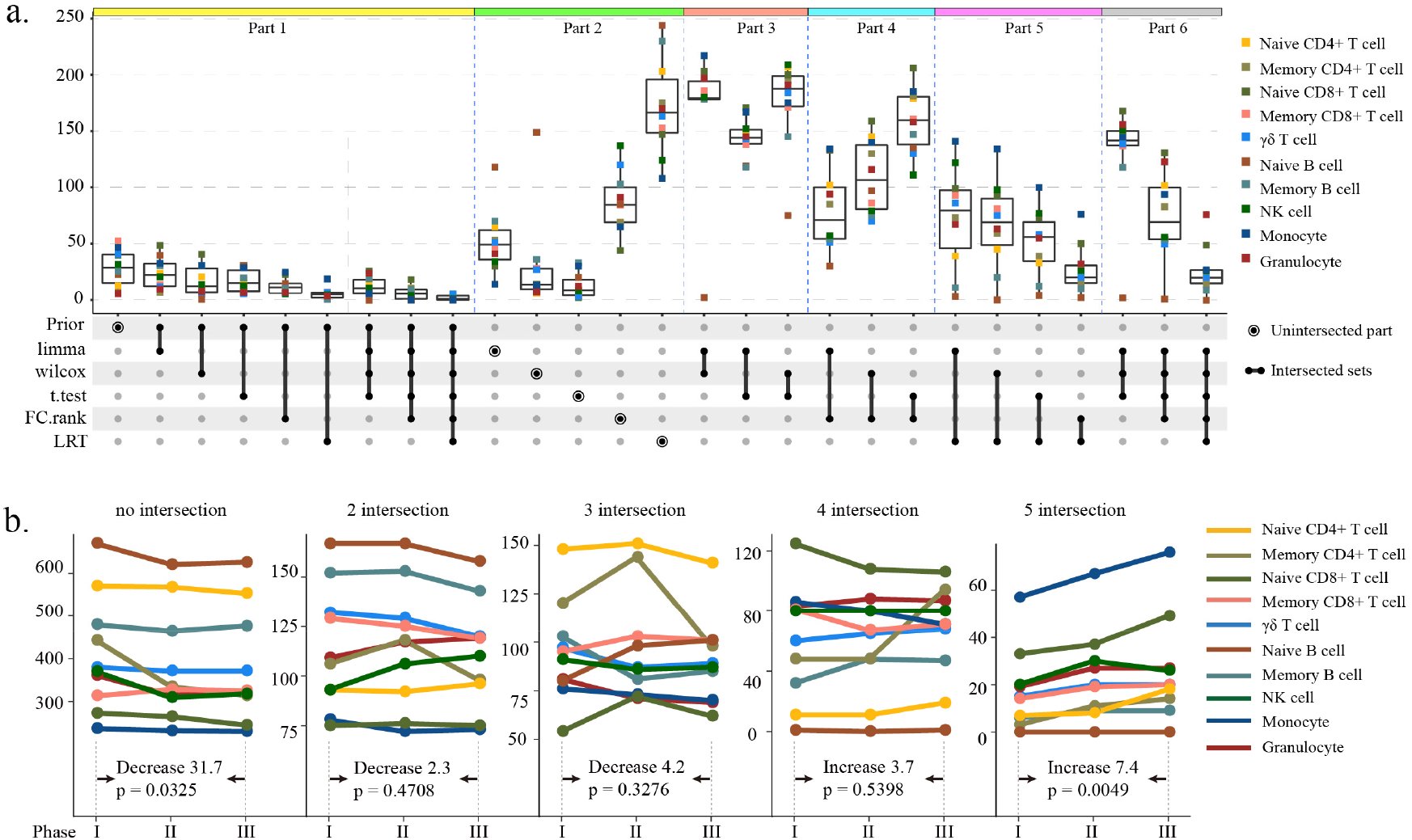
Consistency of specific gene lists obtained by different methods **a**. The intersection of the specific gene lists obtained by the 5 methods and prior knowledge; **b**. The line plot of the intersection number of 5 methods along with the noise cells removal, and in each intersection case, the difference between the interaction on phase I data and phase III data was compared using the t-test, the change amount and the p-value were marked.

### Deep-LDA for single-cell classification and specific gene analysis

By analogizing cell types as topics, and gene proportions as word frequencies, we applied an LDA-based deep generative model on scRNA-seq data analysis (Fig 4a; Methods). The model couples the optimization process of classification with class-specific gene analysis, and directly takes the count matrix as input, which greatly simplifies the analysis process and saves time (Fig 4b, Table S9). We applied the model on phase I data, and finally obtained 24 classes merged from the initial set 50 classes according to the class-specific genes intersection (Fig S6a; Methods). The annotated class distributions were in high concordance with the real cell distribution (Fig 4c). And the inferred specific genes showed high specificity for different classes (Fig 4d; Fig S6b). Then, we re-analyzed class-specific genes by the 5 procedures tested before, and intersected the outcomes with Deep-LDA specific gene lists. The intersection result indicated the specific genes of Deep-LDA had a high degree of consistency with that of the 3 primary methods nominated before (Fig 4d), which suggested Deep-LDA could also be a primary choice for specific gene analysis.

**Figure 4.**
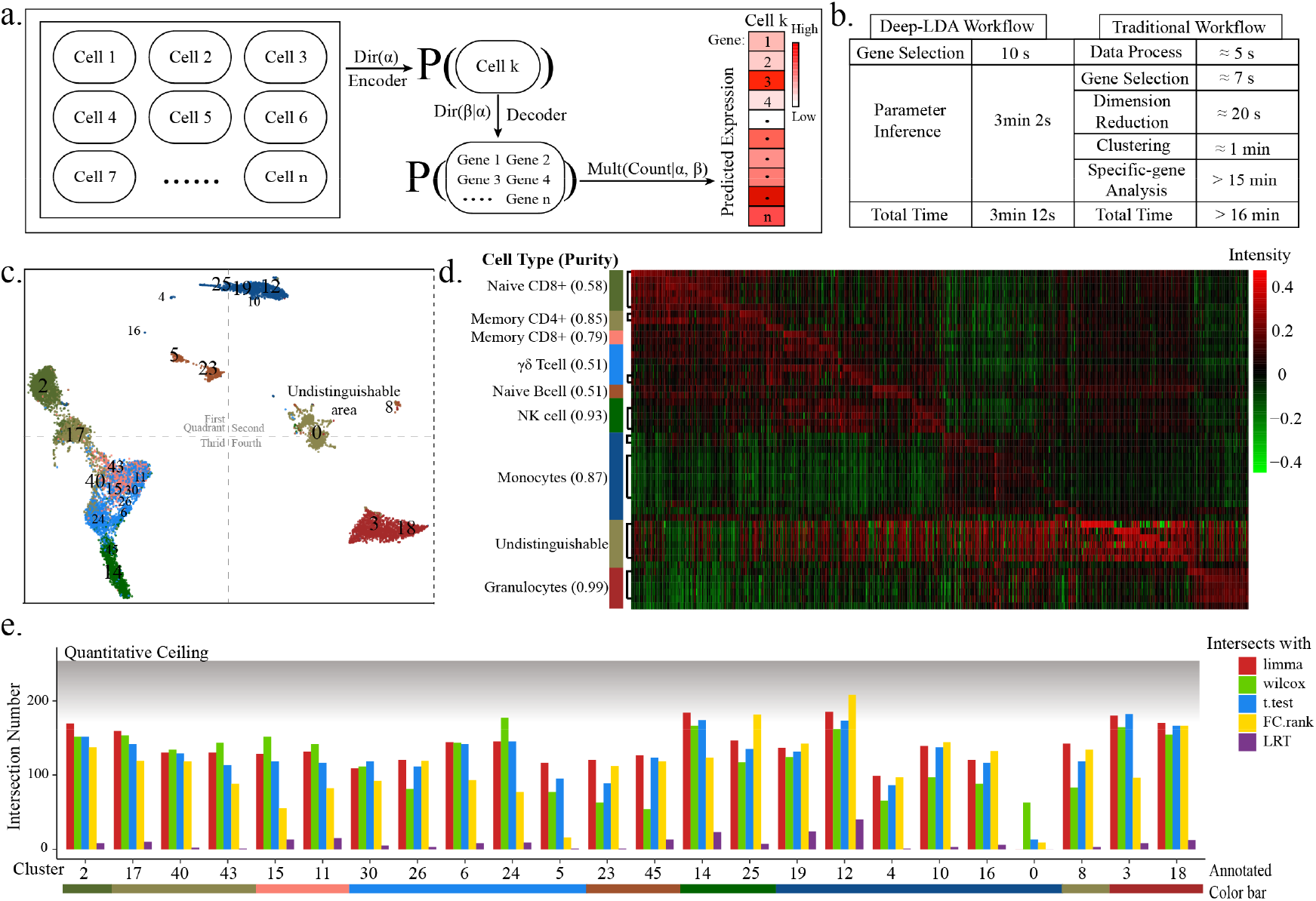
Architecture and Analysis Results of Deep-LDA **a**. Schematic diagram of the model generation process and the corresponding variational model; **b**. Comparison of analytical procedure and time between Deep-LDA and traditional analysis methods; **c**. Clustering result of phase I data. The black numbers marked the position of the cluster centers (cluster index see **Fig S6a**). The clusters were colored by the cell type of the largest proportion, and the cell type color panel is the same as **Fig 1b**; **d**. Heatmap of the β matrix, the class ranking is the same as **Fig S6a**. The left color bar denotes the color of the cluster which is the same as **c**, and the values in parenthesis represent the purity; **e**. The intersections between the specific gene lists of Deep-LDA and that of the other 5 methods, and the quantitative ceiling indicated the maximum intersection number.

### Deep-LDA had outstanding classification performance and lower noise sensitivity

Deep-LDA model was applied on the 3-phase data, whose clustering results had a high consistency with the real distribution at all phases (Fig 5a; Fig S7a). From the annotated memory CD8+ T cell distribution, we found that the distribution shape drawn from this model was more similar with the real distribution shape, and did not form a blocky distribution like other clustering procedures (Fig 5a right, 5b), which suggested Deep-LDA has a higher nonlinear fitting ability. Then, we applied the 13-clustering metrics to compare the Deep-LDA model with the other 5 clustering methods. And the external evaluation score lines of the Deep-LDA model were located above or near the upper lines of other methods’ borders, and exceeded more especially on the phase I part, which indicated Deep-LDA had an outstanding performance and should be one of the first echelon methods (Fig 5c; Fig S7b). In addition, the linear fitted slope of the evaluation score lines revealed that Deep-LDA had lower noise sensitivity than other methods (Fig 5c; Fig S7b). The poor internal clustering scores of Deep-LDA indicated that the outcome of the model was not optimized according to the uniform dimensionality reduction space which was the space for internal clustering metrics calculation, but was optimized according to the inferred feature space of different classes.

**Figure 5.**
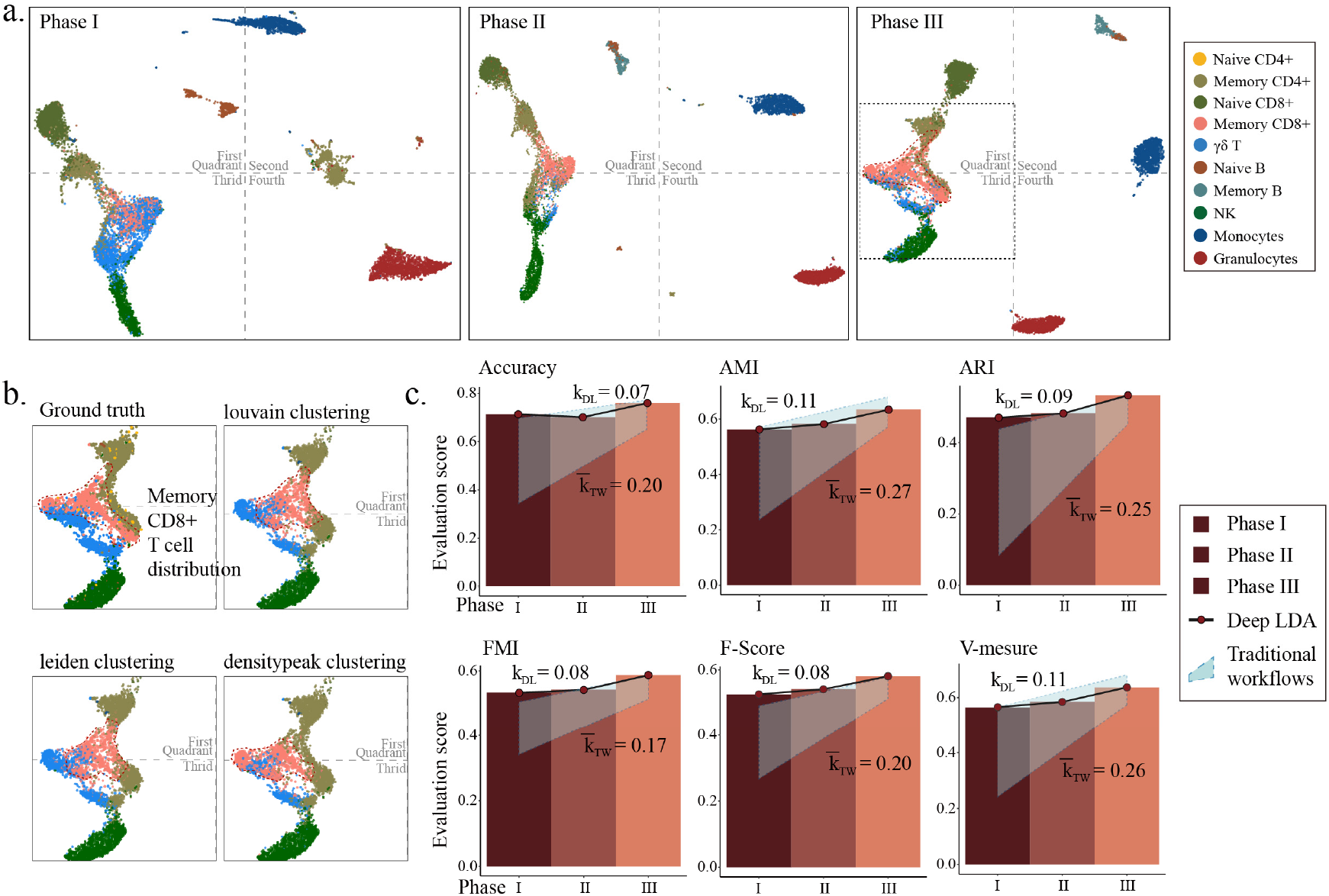
Clustering results of Deep-LDA and the class external evaluation metrics **a**. Deep-LDA clustering results on the three-phase data, and color annotated with the cell type of maximum occupancy cells; **b**. Memory CD8+ T cell distribution area and the same area of the other 3 traditional clustering procedures; **c**. The superposition of the bar and the line plot of the Deep-LDA clustering evaluation score. The light blue quadrilateral box is the outline of the line plots of the other 5 traditional methods, and the upper and lower sides are the fitting lines of the best method and the worst method, respectively. The *K*_*DL*_ is the fitting slope based on the Deep-LDA, and the 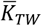 is the average slope of the other 5 methods.

### Single-cell annotation performed by Deep-LDA transfer learning

The encoder in Deep-LDA is for topic assignment, which serves as the cell type classifier. And the decoder is to generate the gene probability distribution, whose parameter was optimized to be class-specific gene frequency (Methods). Thus, we can preload the gene frequencies from independent reference datasets into the decoder, and then freeze the decoder in the parameter inference process, which can generate a single-cell classifier by transferred knowledge. In this project, we used the centroids of the 10 types of cells and an independent immune cell reference dataset^31, 38^ as transferred knowledge (Fig 6a, b left), and then applied the optimized models to perform single-cell annotation (Methods). From the distribution on the UMAP plot, the annotated cell of both inferred results matched well with real related-cell distribution (Fig 6a, b right). After that, we compared the two results with the outcomes from cellID^39^ and singleR^40^ annotation tools with the 13-clustering metrics (Fig S8a, b; Methods), which verified the better performance of Deep-LDA transfer learning (Fig 6c; Fig S8c).

**Figure 6.**
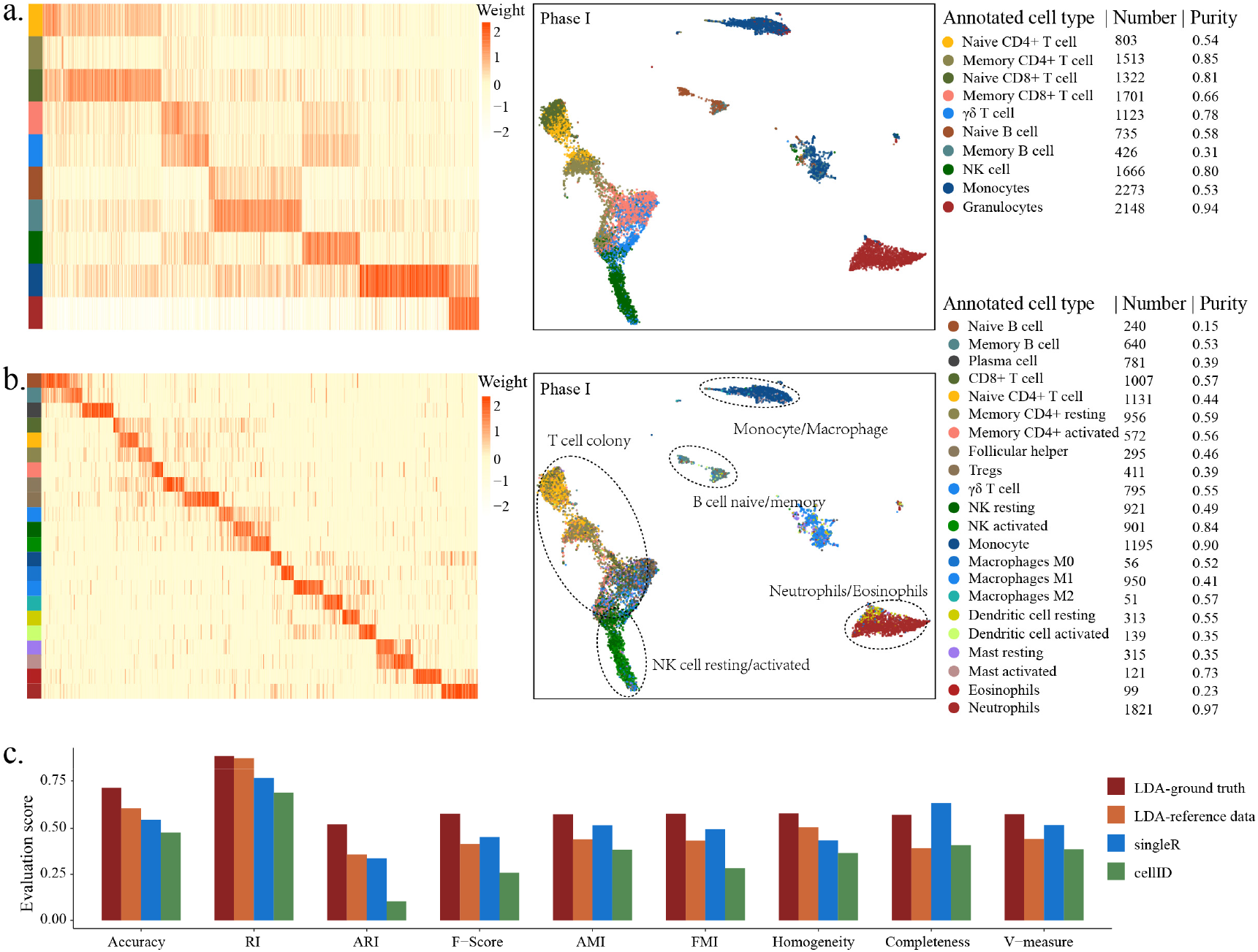
The transfer learning results of Deep-LDA and evaluation metric comparison Deep LDA used **a**. centroids of the 10 types of cells and **b**. an independent reference dataset for transfer learning. The left panel shows the heatmap of pre-obtained β matrix parameters, and the right panel shows the single-cell annotation results, marked with the number and purity for each cell type. **c**. Class evaluation score comparison of Deep-LDA’s annotation results with 2 other annotation methods.

## Discussion

Clustering is the crucial and essential step in the current scRNA-seq data analysis workflow^20^. In this paper, we mainly worked on clustering-related evaluation, optimization and redevelopment. Firstly, we applied scRNA sequencing on a pool of pre-labeled immune cells, by which a batch of data with main and fine labels was generated as ground truth for clustering evaluation. Then, we eliminated two types of noise cells inferred by deep generative models, which improved clustering performance and the consistency of several specific-gene analysis methods. After that, by applying the Deep-LDA model for scRNA-seq data analysis, which couples the classification process with class-specific genes analysis process, that is, the two processes are mutually dependent, we successfully simplified the analysis workflow, shortened the analysis time, and at last verified its superior clustering performance.

The prior knowledge of cell-specific genes is the basis for clustering annotation^20^. However, the recommended reliable specific genes of the current databases were not ranked high and even not significant in our specific lists generated from the pooled sequencing data, and particularly worse in that of the fine-label clusters. More than that, the resolution that the scRNA-seq can achieve was determined by transcriptome features^41^, and it should be much higher than the clusters classified by fine labels, which were defined by cell membrane proteins in our data. So, the current quality and quantity of the specific genes in prior knowledge were insufficient for accurate single-cell annotation^42^. In addition, the quality of the obtained specific gene from most of the current scRNA-seq articles is unstable, because of the unstable clustering quality. Therefore, the more non-randomly captured or pre-labeled single-cells are sequenced, due to the ground truth existence, the more cell-specific gene with higher reliability will be contributed to current databases.

The community-detection algorithm is the primary choice in the current clustering analysis, which was confirmed by the superior performance of louvain- and leiden-based methods in our project. However, an earlier applied method, the density peak clustering, which only grouped cells by the cell density on the two-dimension plot, can basically achieve the same superior performance. The reason is that the community-detected clusters also tend to form distinguishable distribution on UMAP/tSNE, which is the basis for density peak clustering. By specifically surveying the cell distribution on UMAP/tSNE, we found that for the more subdivided cell types, their shapes will be more irregular, and the boundaries between them will be more blurred. We deduced that this undistinguishable distribution will limit the capabilities of both community detection and density peak clustering. The major reason is that the feature space for classes with the different subdivided degrees is treated uniformly, which should have different granularity between inter-main label clusters and inter-fine label clusters. Therefore, we recommend that the initial clusters obtained should be merged into major clusters by similarity index. And then the high variance genes within the major cluster can be re-screened, that is, using finer-grained features of inner-cluster cells, to make a new round of clustering for fine clusters.

The positive contribution of inferred noise cells removal was confirmed in our study. In addition, unlike reshaping expression values for normalization, the excess quality control of noise cells, such as removing the cells in differentiated states, would also have a positive contribution. This deduction should be understood as that we should first map out the class relationships among the more discriminative cells in the rough-grained latent space, and then analyze the relationships of the cells of the less discriminative with these established broad classes. The impurity cell inference model in our project can learn the class affiliation probability. In the result part, the cells with higher impurity probability are always distributed in the junction between two clusters (Fig. S2b). Besides that, the deep generative model is an effective tool for estimating various types of noise cells, which can make a flexible assumption about how the noise cells are involved in the data generation, and then infers the noise probability of each cell by the observation data^27^. In addition to the model inference which has to make an assumption, we found that noise cells had distinguishable distribution patterns on UMAP/tSNE (Fig 2a), so manually selecting abnormally distributed cells should also be considered a good way.

In the Deep-LDA model, the training parameter of specific gene frequency gradually showed different granularity inter- and inner-major clusters, which illustrated that the model is suitable for making classification in the data space of main and fine clusters. And the initial set parameter of the class number describes the resolution for classification. The generative architecture of Deep-LDA in our project was the classical LDA architecture of topic modeling and was not re-designed according to the characteristic of scRNA-seq data, such as incorporating the parameter for controlling the 0-inflation ratio. The neural network structure of the variational model was implemented by fully connected layers. In practice, we had tested CNN, RNN and slide layers in model construction, but they did not show better results. Therefore, by incorporating a more suitable network for learning the transcriptome data characteristics (functional groups and network structure, etc.) and incorporating more regulatory parameters in the generative process, the model will have greater potential in the future.

## Methods

### Participant

A healthy volunteer was enrolled in the study and signed the informed consent of sample collection and data analysis. The study was approved by the Ethics Committee of the Institute of Biomedical Sciences, Fudan University (No. 2018IBSJS014). The 150mL of peripheral blood was extracted by a professional doctor from the subject in the hospital laboratory, and the sample was anticoagulated by EDTA(Ethylene Diamine Tetraacetic Acid) and stored at 2∼8°C.

### Immune cell enrichment

Peripheral blood mononuclear cells (PBMCs) were isolated from the phase above the red blood cells by density gradient centrifugation using LymphoprepTM (STEMCELL Technologies) as per the manufacturer’s instructions. The cells were counted via trypan blue stain and haemocytometer. Then, PBMCs were suspended to 5*10^7 cells/mL. The ten types of immune cells were enriched and purified by EasySep™ Human Series kits based on MicroBeads positive and negative screening (Table S1). The purification kit, surface markers of cells, and heoretical purity are shown in Table S2. The purification process was conducted strictly according to the manufacturer’s instructions. The purified and enriched cells were separated for subsequent single-cell sequencing and bulk sequencing (Figure 1a).

### Pooled sample scRNA-seq

Each type of enriched and purified cell was mixed with a human universal antibody conjugated with Sample Tag (a specific oligonucleotide sequence) for sample identification, which has common 5’ and 3’ ends and the specific ‘Sample Tag sequence’, whose combination is as follows:

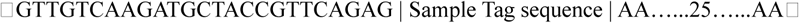

The relationship between cell type and ‘Sample Tag sequence’ is shown in Table S4. After the mixture, 10 types of labeled cells constituted a pooled sample for single-cell sequencing^43^. Single cells were captured by microwell plates and were lysed, and mRNA was hybridized with primer sequences on Cell Capture Beads. The BD Rhapsody™ Targeted mRNA and Abseq Amplification Kit were used for library preparation. The libraries were sequenced on Illumina’s sequencer. Information about Tag and WTA files was found in Table S5.

### Bulk sequencing

10 types of enriched and purified cells were respectively sampled (each sample has more than 100 cells) for bulk sequencing and repeated 3 times. A total of 30 samples were sequenced. The samples were lysed, reverse transcribed into cDNA, and amplified for library preparation. SMART-Seq® HT Kit was used for cDNA amplification. The quality of the cDNA library was evaluated, and 6 samples were excluded. Finally, 24 samples (Table S6) were sequenced on Illumina’s sequencer.

### Specific genes collection

We collected the specific genes from the Cellmarker database^44^, PanglaoDB database^45^, and SC2Disease database^46^. The selected entries for each database are shown in Table S7. The relevant specific genes of immune cell^47^, T cell^48^, CD4+ T cell, CD8+ T cell^49^, and Naive T cell^50^ were obtained from references. The specific genes of Naive T cell were obtained from the intersection of Naive CD4+ T cell and Naive CD8+ T cell. The specific genes of B cell were obtained from the intersection of Naive B cell and Memory B cell. The specific genes appearing in two or more cell types were removed. Finally, a total of 460 specific genes for 16 cell types were collected (Table S8).

### Data processing

Fastq files of the two types of sequencing were quality-checked with FASTQC, and adapters were removed by TrimGalore. For single-cell sequencing data, Cell barcode and UMI sequence were estimated and extracted by UMItools in the R1 files, and were then inserted into the R2 files. The R2 files were processed by STAR for single-end alignment. As for bulk-seq sequencing data, the alignment was paired-end.

The reference genome was GRCh38(www.ncbi.nlm.nih.gov/assembly/GCF_000001405.39). The annotation was Genecode v.29(www.gencodegenes.org/human/release_29.html). Read count was adjusted by the correspondent UMIs. The true relationship between cell barcode and Sample Tag Sequence was estimated via Seurat’s Demultiplexing HTOs function (Demultiplexing with hashtag oligos) in Tag files^43^.

Then, the count matrix was log-transformed. 2000 genes with the maximum variance were selected and scaled. The top 30 principal components (PCs) were obtained by PCA dimensional reduction. And the cell distribution was visualized by UMAP and tSNE based on those PCs. For the 3-phase data with a stepwise noise cells removal, the data processing was the same.

### The composition of bulk-seq data

The 2000 genes from the data processing were used as gene signatures for deconvolution. And then, the centroids of 10 types of immune cells were calculated to be the Signature gene file. Then, we used CIBERSORT software to decompose the bulk-seq data by the signature gene file and to verify the consistency between scRNA-seq data and bulk-seq data.

### Clustering Analysis

A total of five algorithms, densitypeak clustering, DDRTree (Discriminative dimensionality reduction via learning a tree) clustering, louvain clustering, leiden clustering, and consensus clustering, were used for clustering. Densitypeak clustering and DDRTree clustering were implemented by monocle (version 2.22.0). The tSNE coordinate was used for densitypeak clustering. The DDRTree dimension-reduced data was used for DDRTree clustering. The clustering was implemented by the clusterCell function of monocle with ‘method’ equal to ‘densityPeak’ or ‘DDRTree’. Louvain clustering and leiden clustering were implemented by monocle3 (version 1.0.0) with PCs (dim=30), the cluster_cells function with ‘method’ equal to ‘louvian’ or ‘leiden’. Consensus clustering was implemented by the scena_cpu function of SCENA(version 1.0.3). The cluster number was set as 18, which led to the optimal results.

### Specific genes analysis

The count matrix was normalized as recommended. The pairwise comparison strategy was used for specific genes analysis, where the core differential analysis algorithms were by t-test, limma, wilcox, logFC ranking, and LRT (likelihood ratio test). The analysis based on the t-test was implemented by the findMarkers function of scran (version 1.22.1). Limma used the one-VS-others pairwise strategy for all clusters. The analysis based on wilcox was implemented by the FindAllMarkers function of Seurat (version 4.0.5). The analysis based on LogFC ranking was implemented by the getClassicMarkers function of singleR (version 1.8.1). The analysis based on the LRT was implemented by the differentialGeneTest function of monocle (version 2.22.0). Then, for the differential gene lists of each cell type obtained from the different analyses, the 250 top-ranking genes were selected as specific genes.

### Cell Type Automatic Annotation

SingleR^40^ (version 1.8.1) and CelliD^39^ (version 1.4.0) were used for cell type automatic annotation. The reference data with known cell types was used to annotate experimental data, where HumanPrimaryCellAtlas and PanglaoDB_markers_27_Mar_2020 were, respectively, the reference data of SingleR and CelliD.

### Intra-cluster Evaluation Index

Silhouette Coefficient index was a measure of the cohesion and separation of clusters and tended to be higher, when the distance between samples of the same type was smaller and between samples of the different types was larger. The formula was the following:

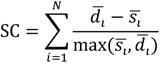

Where N denoted the number of all samples. *s*_*i*_ denoted the average distance between sample i and all the other samples with the same cell type label. *d*_*i*_ denoted the average distance between sample i and all the other samples with the nearest and different sample type labels. The score was evaluated via metrics.silhouette_score function of sklearn.

Calinski-Harabaz index is a measure of cohesion via covariance. The intra-cluster is closer and the inter-cluster is more scattered, so the score is higher. The formula was the following:

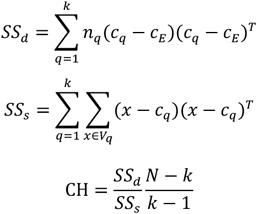

Where N denoted the number of all samples, k denoted the number of clusters, *n*_*q*_ was the number of samples within cluster q, *c*_*E*_ was the centroid of all samples, *V*_q_ was the set of cluster q, *SS*_*d*_ was the sum of the covariance of each cluster centroid, *SS*_*s*_ was the sum of covariance of the sample within the same cluster. The value was calculated by metrics.calinski_harabaz_score function of sklearn.

Compactness index was the average distance between the data point and the centroid of the cluster. The formula was the following:

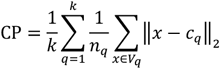

Where the meaning of parameters unexplained was identical with those described above.

Davies-Bouldin index was to calculate the averaged inner-cluster distance of 2 clusters, which was divided by the distance of the centroids of 2 clusters, and then found the maximum. The score tended to be lower when the inner-cluster distance was smaller, and the intra-cluster distance was larger. The formula was the following:

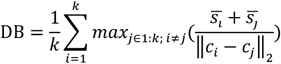

Where the meaning of parameters unexplained was identical with those described above. The value was calculated by metrics davies_bouldin_score of sklearn.

### Inter-cluster Evaluation Index

Purity was calculated as the following formula:

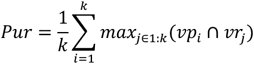

Where *vp*_*i*_ was the sample set of clusters i, *vr*_*j*_ was the sample set of clusters j, and to obtain the maximum number of intersections between *vp*_*i*_ and *vr*_*j*_.

Rand index was the ratio of true prediction and was calculated as the following formula:

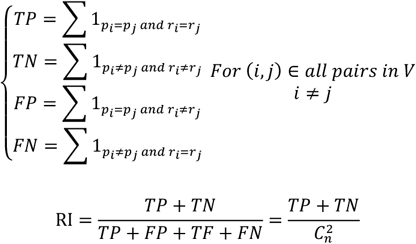

Where V was the set of all samples, *p*_*i*_ was the predicted cell type of sample i, *r*_*i*_ was the true cell type of sample i, n was the sample number, 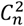 was the combination calculation.

Adjusted Rand index was the standardized Rand index, and the formula was the following:

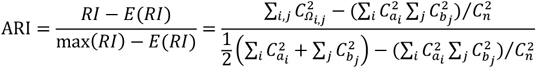

Where *Ω*_*i*,*j*_ was the intersection number of predicted cluster i and true cluster j. *a*_i_ was ∑_*j*_ *Ω*_*i*,*j*_ and *b*_*j*_ was ∑_*i*_ *Ω*_*i*,*j*_. The value was calculated by metrics.adjusted_rand_score of sklearn.

F-score was defined as:

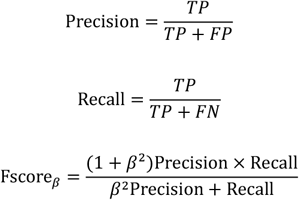

Where the meaning of parameters unexplained was identical with those described above. β was the hyperparameter and was set to 1.

Fowlkes-Mallows index was the geometric mean of pair-wise precision and recall.

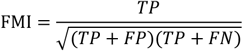

Where the meaning of parameters unexplained was identical with those described above. The value was calculated by metrics.fowlkes_mallows_score function of sklearn.

AMI (Adjusted mutual information) index:

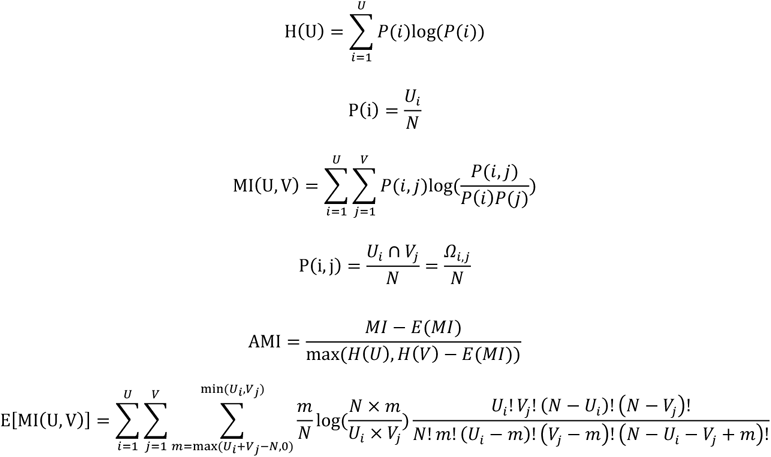

Where the meaning of parameters unexplained was identical with those described above. U and V represented the predicted cluster and true cluster, P(i) was the proportion of cells labeled i., P(i, j) was the proportion of the cells of predicted label i and true label j. The value was calculated by metrics.adjusted_mutual_info_score of sklearn.

Homogeneity measured whether all clusters fell into the same clusters with the true label. The formula was the following:

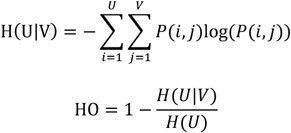

Where the meaning of parameters unexplained was identical with those described above. The index was calculated by metrics.homogeneity_score of sklearn.

Completeness measured the extent to which samples belonging to the true same cluster fell into the predicted same cluster. The formula was the following:

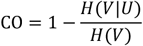

The value was calculated by metrics.completeness_score of sklearn.

V-measure was the harmonic mean between homogeneity and completeness. The formula was as follows:

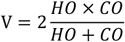

The value was calculated by metrics.v_measure_score function of sklearn.

### Pseudo cell generative model

The model was constructed according to the structure of cellbender^27^, and the observed data *X*_*ng*_ was generated as follows:

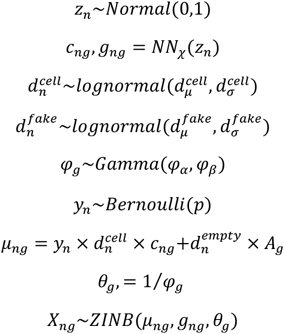

Where *z*_*n*_ was the latent variable, which was hypothetically generated according to the normal distribution. During the inference process, an encoder (encoder_z) was used as a variational model with the input of gene counts for generating the mean and variance of *z*_*n*_. *c*_*ng*_ was the generated true gene expression values derived from latent space through an decoder (*NN*_*χ*_*). g*_*ng*_ was the probability that gene was not observed. 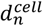 was the scale factor of the true cell, and 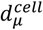 was generated by encoder_d in the inference process. 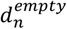 was the scale factor of the pseudo cell. *y*_*n*_ was the label of whether the cell was a true cell, where the probability was generated by endoder_p in the inference process. *µ*_*ng*_ was the generated true gene expression values after combining the hypothesis of the pseudo cell, where *A*_*g*_ was the ambient RNA. *φ*_*g*_ was inverse_dispersion parameter of the model. The final observation value *X*_*ng*_ was generated by ZINB (Zero-inflated Negative Binomial) model. The parameter priors of the generation process are given in Table S10.

### Impurity cell generative model

There exited impurity cells in each cell cluster, because the cells’ purification degree was less than 100% (Table S2). The class affinity probability for each cell was inferred by the semi-supervised generative model^51^ proposed by scANVI^52^. The observed data *X*_*ng*_ was generated as following:

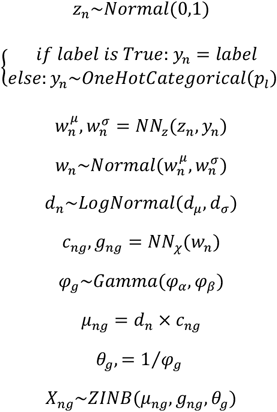

Where the meaning of *z*_*n*_, *c*_*ng*_, *g*_*ng*_, *µ*_*ng*_ and *φ*_*g*_ was the same as in the Pseudo Cells Inference Model. *w*_n_ was generated by a decoder with latent variable *z*_*n*_ and cell label *y*_*n*_ as input. Because Impurity Cells Inference Model was a semi-supervised model, we set 50% of cell labels were unknown. In the unknown case, the cell label *y*_*n*_ was sampled from *OneHotCategorical* distribution and the parameter *p*_*l*_ was the probability of the cell label generated by encoder_p in inference process. The parameter priors of the generation process are given in Table S10.

### Generative model inference

The posterior parameter distribution inference was combined with the prior distribution and observed data X. And because the p(λ_1_ … λ_*k*_|*X*_*n*_) can hardly be deduced, we used the variation inference to infer the approximately posterior distribution. Variation distribution can be divided into independent distributions of different parameters. The formula was as follows:

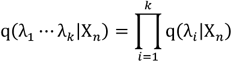

The likelihood function was deduced as:

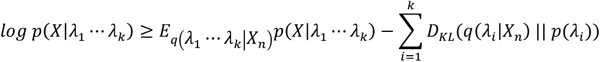

The evidence lower bound (ELBO) was set as the optimization objective. The optimization was realized by pyro. The loss models and the optimizer for the three models described above were ELBO and Adam.

### Gene selection for the deep generative model

Genes that are expressed or not expressed in the vast majority of cells are more useless in distinguishing cell types, so we select genes expressed in a certain percentage of cells for the optimization of the deep generative model. This project used genes expressed in cells that account for 40∼90%, a total of 2441 genes.

### Deep-LDA model

This model is an LDA-based (Latent Dirichlet Allocation) deep generative model. And the generation structure was referenced from proLDA^30^. The gene frequency was analogized as word frequency, and the cell type distribution was analogized as the topic distribution. The observed data *X*_*ng*_ was as follows:

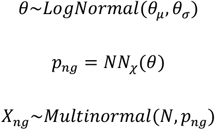

Where *θ* denoted the cell type distribution, which should be generated by the Dirichlet distribution and here generated by a lognormal distribution. In the inference process, *θ* was generated by an encoder (variation model) with the input of the gene counts. *p*_*ng*_ denoted the gene probability distribution, which was generated by a decoder (*NN*_*χ*_) and could be represented as *p*_*ng*_ = *softmax*(*β* × *θ*). So, the weight *β* can be used as the training result of gene frequency for different topics or cell types. The observation value *X*_*ng*_ was generated by the *Multinormal* distribution and N was all genes count of a cell. The parameter priors of the generation process are given in Table S10.

### Specific gene of Deep-LDA

The optimized *β* matrix (*N*_class_ x *N*_gene_) in the generative model was used as the basis for specific gene screening. That is, for the optimized gene frequency list of each class, a certain number (default is 250) of high-frequency genes were selected as specific genes for the class.

### Class merging of Deep-LDA

Based on the specific gene intersections of different classes obtained by Deep-LDA, a threshold was set and classes with the number of intersected genes above this threshold were merged. In this process, we suggest to select more specific genes for each class (for phase I ∼ phase III data, the number of specific genes used for merging classes are set as: 1700, 1400, 600), and set a certain threshold (for phase I ∼ phase III data, the thresholds for merging are: 1500, 1000, 300) to produce the merged cluster number as expected (for phase I ∼ phase III data, the number of clusters are: 24, 18, 21).

### Transfer learning of Deep-LDA

We first intersected the gene list of the signature matrix (transferred knowledge) with that of the data to be analyzed so that both matrixes contained the same gene signatures. Here, we used two types of signature files. The first one being the centroid of each cell type through our pooled sample sequencing data, and the 2441 genes for the deep generative model served as gene features. The second one is obtained independently from the human genome array (GSE22886)^38^, containing a total of 22 cell types, and the gene features are selected according to the 530 specific genes supplied by CIBERSORT^31^. Then, the signature matrix (count or raw array intensity data type) was scaled and loaded into the β parameter of the decoder. Deep-LDA was then performed on inference with the data to be analyzed while the decoder parameters were fixed. The encoder obtained after optimization was used as a classifier for annotating new data using transferring knowledge.

## Data availability

The project contains a total of 1 pooled sample scRNA-seq data and 24 bulk sequencing data, which have been uploaded to SRA. The samples and uploaded data information are shown in https://dataview.ncbi.nlm.nih.gov/object/PRJNA843175?reviewer=s4iefbplretvm0005bsj2toc5o. The data will be available with the publication of this article.

## Code availability

The three deep generative models covered in this article, as well as necessary clustering evaluation tools and plotting tools are organized into the ScDeepTools (https://github.com/liuyunho/ScDeepTools) software. In addition, the software includes an interactive cell distribution plotting tool (**Fig S9**) for manually cell picking and deletion.

## Author contributions

Conceptualization, Yunhe Liu, Gang Liu and Lei Liu; experiment, Yunhe Liu, Qiqing Fu, Chenyu Dong, Xiaoqiong Xia; methodology, Yunhe Liu, Qiqing Fu; software, Yunhe Liu; validation, Yunhe Liu, Qiqing Fu; manuscript preparation, Yunhe Liu, Qiqing Fu, Chenyu Dong; visualization, Yunhe Liu, Qiqing Fu, Chenyu Dong; funding acquisition, Lei Liu. All authors have read and agreed to the published version of the manuscript.

## Funder

L.L. was supported by the grant No. 2021YFC2500403 from the National Key Research and Development Program of China.

## Acknowledgements

For technical advice and financial assisting, we thank the project of Shanghai’s Double First-Class University Construction and Development of High-Level Local Universities: Intelligent Medicine Emerging Interdisciplinary Cultivation Project, and also Medical Research Data Center of Fudan University.

## Notes

### Competing Interest Statement

The authors have declared no competing interest.

