## Supplementary Table S1--S7, S9-S10 for "Optimization and redevelopment of single-cell data analysis workflow based on deep generative models"

**Supplement Table 1-7, 9-10:**

**Table S1** Information of the expected cell enrichment experiments for each cell type

| Cell type | Cell content in the whole blood（10^6 cells/mL） | The estimated amount of blood required（mL） | Theoretical cell amount（10^6 cells） | The estimated amount of cells to be obtained at a recovery rate of 20%-50%（10^4 cells） |
|---|---|---|---|---|
| Naive CD4+ T Cells | 0.08 – 0.76 | 5* | 0.4 - 3.8 | 8 - 190 |
| Memory CD4+ T Cells | 0.25 – 0.81 | 12.5* | 3.1 - 10 | 62 - 500 |
| Naive CD8+ T Cells | 0.03 – 0.21 | 12.5* | 0.37 - 2.6 | 7 - 130 |
| Memory CD8+ T Cells | NA | 10* | 0.08 - 0.72 | 1.6 - 36 |
| Gamma/Delta T Cells | NA | 25* | NA | NA |
| Naive B cells | 0.05 – 0.37 | 50* | 2.5 - 18.5 | 50 - 920 |
| Memory B cells | NA | NA | 0.7 - 6 | 14 - 300 |
| NK Cells | 0.08 – 0.43 | 12.5* | NA | NA |
| Monocyte Cells | 0.20 – 0.90 | 12.5* | 2.5 - 11.2 | 50 - 560 |
| Granulocyte Cells | 2.13 – 6.35 | 0.5 | 1 - 3 | 20 - 150 |

**Table S2** Information of immune cell enrichment kits, cell surface markers, and theoretical purity.

| Cell type | Catalog and kit | Cell surface markers | Theoretical purity |
|---|---|---|---|
| Naive CD4+ T Cells | 17555 EasySep™ Human Naive CD4+ T Cell Isolation Kit II | CD3+CD4+CD45RA+CD45RO- | 96.6 ± 1.5% |
| Memory CD4+ T Cells | 19157 EasySep™ Human Memory CD4+ T Cell Enrichment Kit | CD4+CD45RA-CD45RO+ | 86 - 98% |
| Naive CD8+ T Cells | 17968 EasySep™ Human Naive CD8+ T Cell Isolation Kit II | CD8+CD45RA+CCR7+ and CD45RO-CD57-CD56- | 93.7± 2.4% |
| Memory CD8+ T Cells | 19159 EasySep™ Human Memory CD8+ T Cell Enrichment Kit | CD8+CD45RA-CD45RO+ | 72 - 92% |
| Gamma/Delta T Cells | 19255 EasySep™ Human Gamma/Delta T Cell Isolation Kit | TCR gamma/delta+CD3+ | 90 - 97% |
| Naive B Cells | 17864 EasySep™ Human Memory B Cell Isolation Kit | CD3-CD19+CD27- | 93 ± 5% |
| Memory B Cells | | CD19+CD27+ | 97 ± 2% |
| NK Cells | 17955 EasySep™ Human NK Cell Isolation Kit | CD3-CD56+ | 85.0 ± 8.0% |
| Monocyte Cells | 19359 EasySep™ Human Monocyte Isolation Kit | CD14+CD45+ | 89.7± 3.4% |
| Granulocyte Cells | 19659 EasySep™ Direct Human Pan-Granulocyte Isolation Kit | granulocytes（neutrophil [CD66b+CD16+], eosinophil [CD66b+CD16-] and basophil [CD66b-CD123+]） | 98.4± 1.5% |

**Table S3** Information of immune cells after the experiment of enrichment and purification

| Sample name | Total cell counts | Cell survival rate (%) | Cell concentration (cells/μL) |
|---|---|---|---|
| Naive CD4+ T Cells | 10000 | 86 | 180 |
| Memory CD4+ T Cells | 11000 | 87 | 190 |
| Naive CD8+ T Cells | 9300 | 93 | 155 |
| Memory CD8+ T Cells | 35000 | 86 | 590 |
| Gamma/Delta T Cells | 3300 | 83 | 55 |
| Naive B cells | 5300 | 96 | 350 |
| Memory B cells | 6000 | 90 | 100 |
| NK Cells | 5400 | 90 | 90 |
| Monocyte Cells | 635000 | 92 | 2540 |
| Granulocyte Cells | 900000 | 88 | 3600 |

Description: Cell survival rate > 80%; Without cell adhesion (the conglobation rate < 10%); Without cell fragments or other particles larger than 40μm.

**Table S4** Correspondence between cell type and sample tag sequence.

| Tag name | Cell type | Sample Tag sequence |
|---|---|---|
| Tag 1 | Naive CD4+ T Cells | ATTCAAGGGCAGCCGCGTCACGAT TGGATACGACTGTTGGACCGG |
| Tag 2 | Memory CD4+ T Cells | TGGATGGGATAAGTGCGTGATGGA CCGAAGGGACCTCGTGGCCGG |
| Tag 3 | Memory B Cells | CGGCTCGTGCTGCGTCGTCTCAAG TCCAGAAACTCCGTGTATCCT |
| Tag 4 | NK Cells | ATTGGGAGGCTTTCGTACCGCTGC CGCCACCAGGTGATACCCGCT |
| Tag 5 | Monocyte Cells | CTCCCTGGTGTTCAATACCCGATGT GGTGGGCAGAATGTGGCTGG |
| Tag 6 | Naive B Cells | TTACCCGCAGGAAGACGTATACCC CTCGTGCCAGGCGACCAATGC |
| Tag 7 | Naive CD8+ T Cells | TGTCTACGTCGGACCGCAAGAAGT GAGTCAGAGGCTGCACGCTGT |
| Tag 8 | Memory CD8+ T Cells | CCCCACCAGGTTGCTTTGTCGG ACGAGCCCGCACAGCGCTAGGAT |
| Tag 9 | Granulocyte Cells | GTGATCCGCGCAGGCACACATAC CGACTCAGATGGGTTGTCCAGG |
| Tag 10 | Gamma/Delta T Cells | GCAGCCGGCGTCGTACGAGGCAC AGCGGAGACTAGATGAGGCCCC |

**Table S5** Information of sequencing library and raw data for the pooled-sample sequencing

| Sample name | Concentration (ng/μL) | Main peak length (bp) | Raw data size |
|---|---|---|---|
| Tag | 2.36 | 286 | R1(4.35GB)/R2(3.58GB) |
| WTA | 13.9 | 636 | R1(56.03GB)/R2(60.42GB) |

Description: Tag：specific oligonucleotide tag sequencing files；WTA：sample gene expression sequencing files.

**Table S6** Library information of each cell type in bulk sequencing

| Sample name | Concentration (ng/μL) | Main peak length (bp) |
|---|---|---|
| Naive CD4+ T cell sample 1/2/3 | 19.7/13.3/10.8 | 304/320/323 |
| Memory CD4+ T cell sample 1/2/3 | 14.7/21/16.4 | 305/273/283 |
| Memory B cell sample 1/2/3 | 21.2/21.4/16.7 | 288/288/316 |
| NK cell sample 1/2/3 | 23.4/24.4/20.6 | 299/300/300 |
| Naive B cell sample 1/2/3 | 18.7/20.4/23.4 | 304/299/291 |
| Naive CD8+ T cell sample 1/2/3 | 25.2/19.5/20.2 | 301/293/291 |
| Memory CD8+ T cell sample 1/2/3 | 19.4/21/14.2 | 292/289/288 |
| Gamma delta T cell sample 1/2/3 | 24.6/18.1/23.2 | 294/315/293 |

**Table S7** Information of selected cell entries in the corresponding database.

| Cell type | Database | The selected cell entries |
|---|---|---|
| Naive CD4+ T Cells | Cellmarker | Naive CD4+ T cell |
| Memory CD4+ T Cells | Cellmarker | CD4+ memory T cell， Effector CD4+ memory T (Tem) cell，Effector CD8+ memory T (Tem) cell |
| | SC2Disease | Effector memory CD4+ T cells sort by disease desc.csv |
| Naive CD8+ T Cells | Cellmarker | Naive CD8+ T cell |
| Memory CD8+ T Cells | Cellmarker | Effector CD8+ memory T (Tem) cell |
| γδT Cells | panlaodb | Gamma delta T cells |
| Naive B Cells | panlaodb | B cells naive |
| Memory B cell | Cellmarker | Class-switched memory B cell, Non-switched memory B cell, Memory B cell, Switched memory B cell, Double-negative memory B cell |
| | panlaodb | B cells memory |
| Monocytes | Cellmarker | Classical monocyte, Intermediate monocyte , Non-classical monocyte, Suppressive monocyte, CD14+CD16+ monocyte |
| | panlaodb | Monocytes |
| Granulocytes | Cellmarker | Granulocyte, Neutrophil |
| NK cell | Cellmarker | Natural killer cell, pro-Natural killer cell |
| | panlaodb | NK cells |

**Table S9** Run time of each clustering algorithm

| | LDA | | Louvain | Leiden | densityPeak | DDRTree | Consensus |
|---|---|---|---|---|---|---|---|
| Gene Selection | 10s | Data process | 10.52 s | 10.52 s | 4.75 s | 4.75 s | 39.62 s |
| Parameter inference | 3min 2s | Dimension reduction | 6.29 s | 6.29 s | 3min22s | 2h10min 49s | \ |
| | | Clustering | 10.70s | 10.84s | 3min36s | 2h14min 55s | 21h7min51s |
| | | Specific-gene Analysis | 45min15s | 45min 15s | 3h29min 6s | 3h29min 6s | \ |
| Total time | 3min 12s | Total time | 45min43s | 45min 43s | 3h36min 9s | 7h54min 55s | 21h8min31s |

**Table S10** The parameter prior for generative models used in our project

| Pseudo cell generative model | | Impurity cell generative model | | Deep-LDA model | |
|---|---|---|---|---|---|
| Parameters | Prior | Parameters | Prior | Parameters | Prior |
| $\mu\ for\ z_n$ | 0.0 | $\mu\ for\ z_n$ | 0.0 | $\theta_\mu$ | 0.0 |
| $\sigma\ for\ z_n$ | 1.0 | $\sigma\ for\ z_n$ | 1.0 | $\theta_\sigma$ | 1.0 |
| $d_\mu^{cell}$ | 8.0 | $p_i$ | 0.5 | \ | \ |
| $d_\sigma^{cell}$ | 0.2 | $d_\mu$ | 8.0 | \ | \ |
| $d_\mu^{fake}$ | 8.0 | $d_\sigma$ | 0.2 | \ | \ |
| $d_\sigma^{fake}$ | 0.2 | $\varphi_\alpha$ | 0.2 | \ | \ |
| $\varphi_\alpha$ | 0.2 | $\varphi_\beta$ | 0.2 | \ | \ |
| $\varphi_\beta$ | 0.2 | \ | \ | \ | \ |
| $\log(p)$ | 2.2 | \ | \ | \ | \ |
