## Supplementary Figure S1-S9 for "Optimization and redevelopment of single-cell data analysis workflow based on deep generative models"

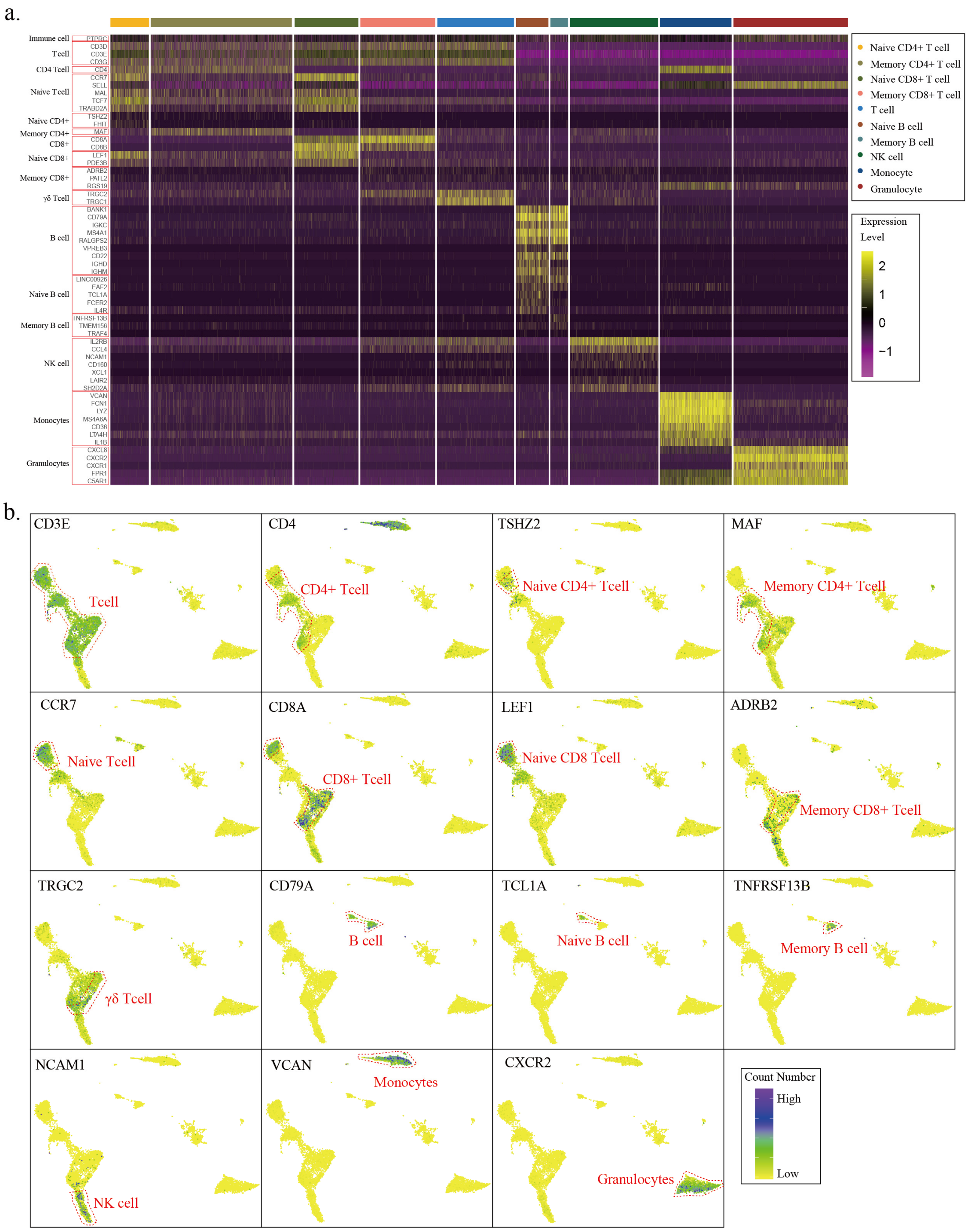


**Figure S1** Gene expression distribution in pooled sample scRNA-seq data

**a.** Robust specific-gene heatmap for pooled scRNA-seq data; **b.** Distribution of the intensity of key specific genes and the red dotted line is used to mark the specific cell distribution boundary.

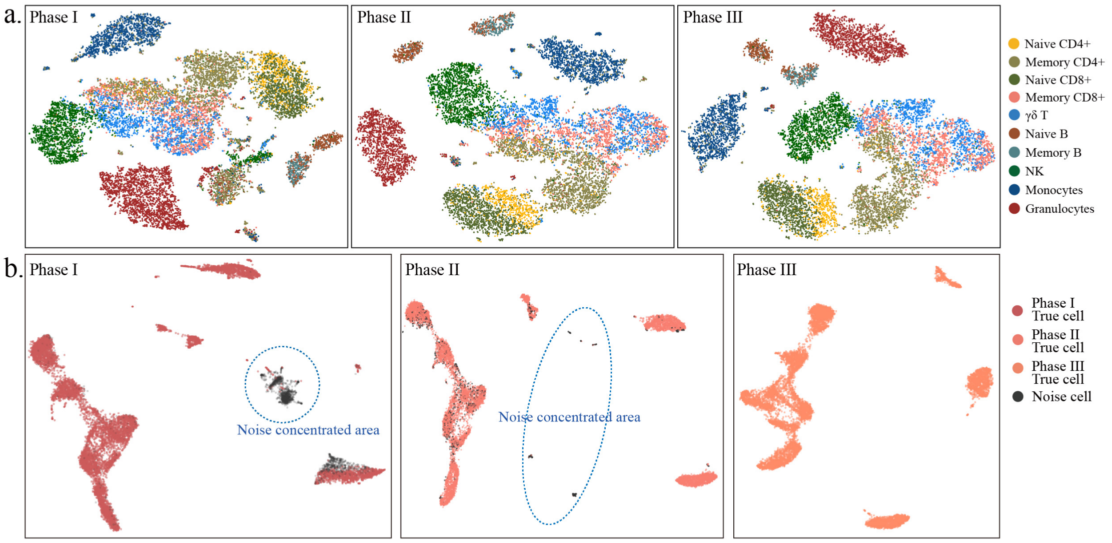


**Figure S2** Noise probability estimation and TSNE visualization after noise removal

**a.** TSNE visualization of the three-phase data, color-coded labels for pre-enriched cell types; **b.** Estimated noise cell probability distribution, including the estimated probability of pseudo-cells in Phase I (left panel), and the estimated probability of impurity cells in Phase II (middle panel).

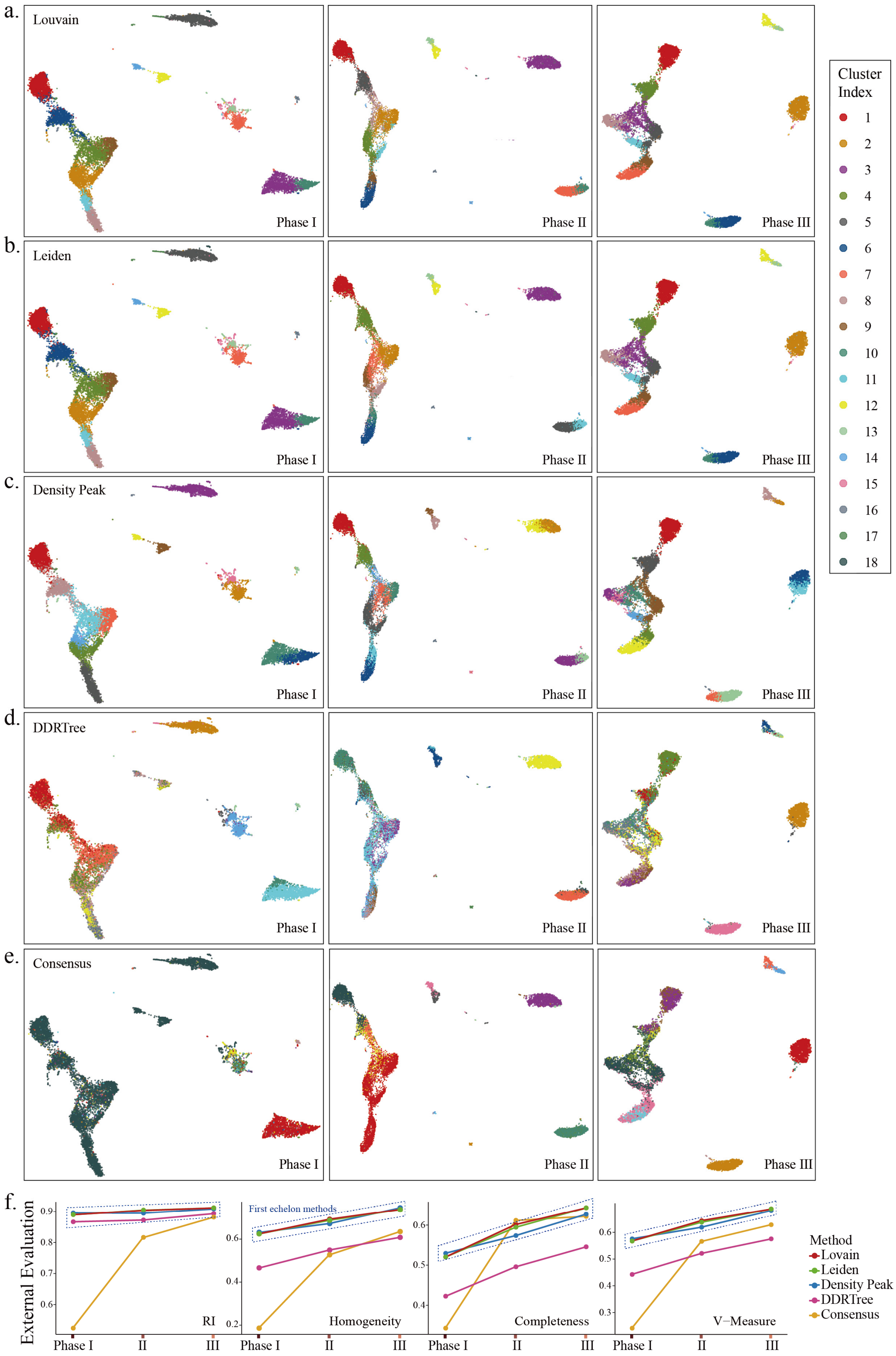


**Figure S3** Clustering of 5 traditional procedures and their evaluation results

The figure contains the results of 5 traditional clustering methods on the 3-phase data (**a-e**) and the line plot of 4 class external evaluation metrics (**f**).

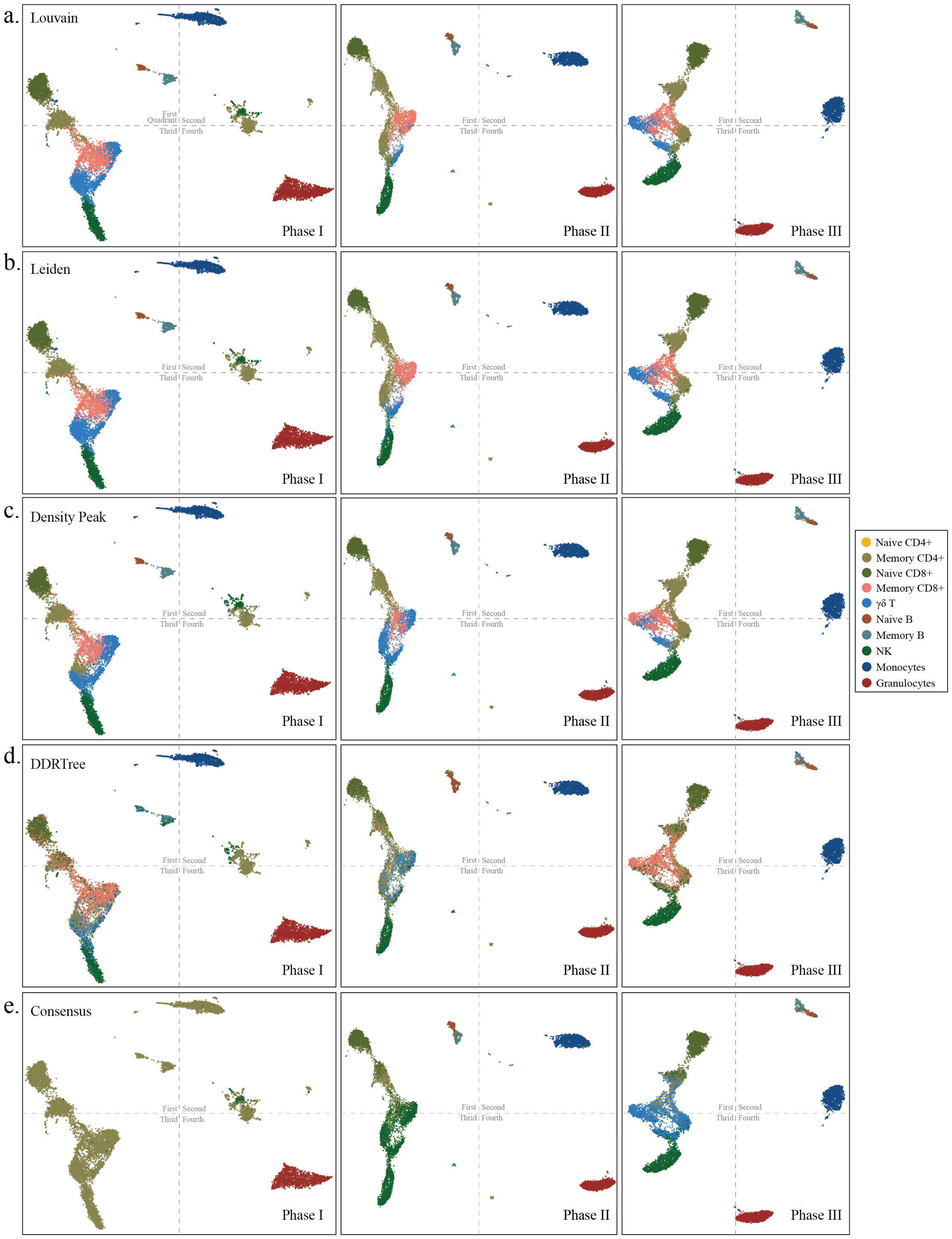


**Figure S4** Cluster annotations for 5 traditional clustering procedures

The clustering results of the five traditional clustering processes are annotated using the class maximum occupancy cells (**a-e**) and labeled with the same color scheme as Fig. 1b, Fig 2a

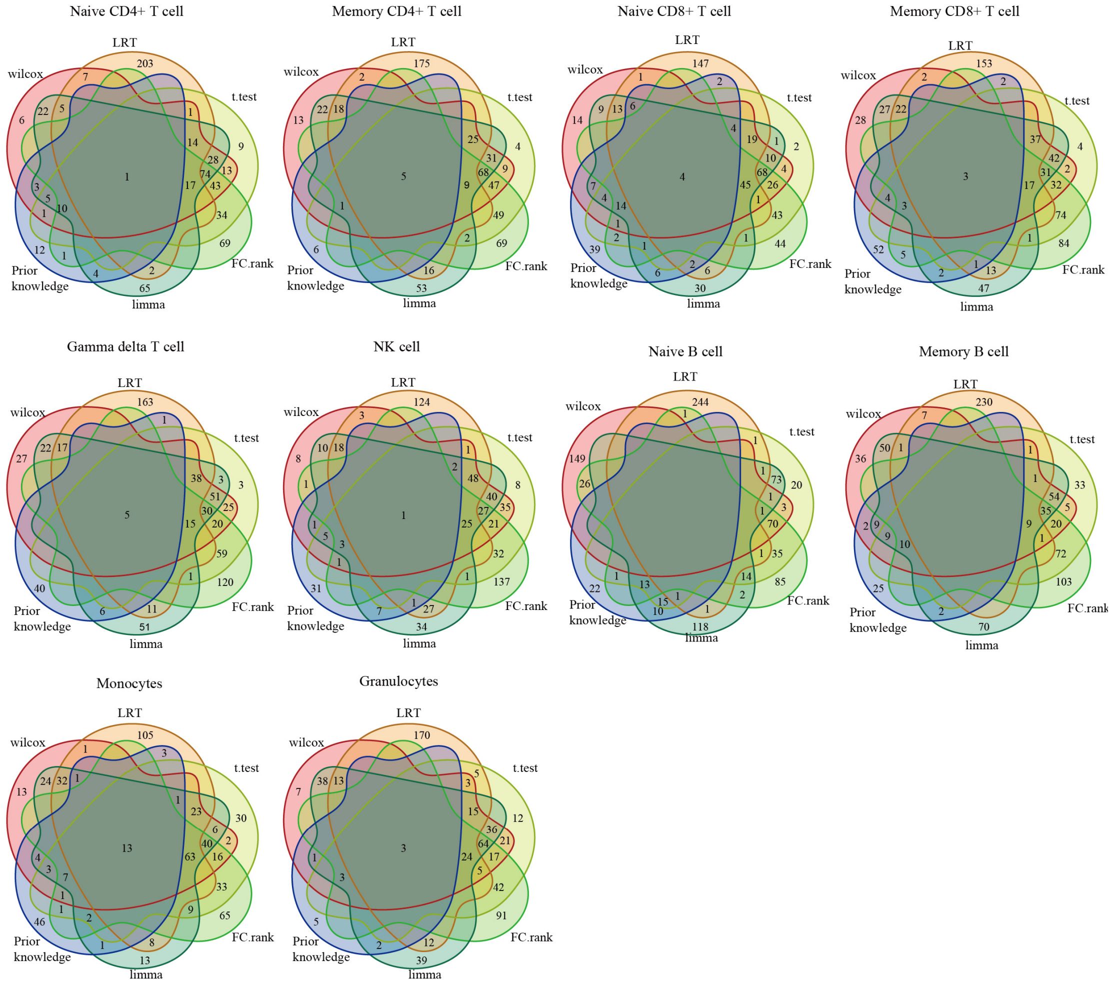


**Figure S5** Intersection of the class-specific gene lists

This figure shows the intersection Venn plot of the class-specific genes lists obtained by the 5 methods and prior knowledge.

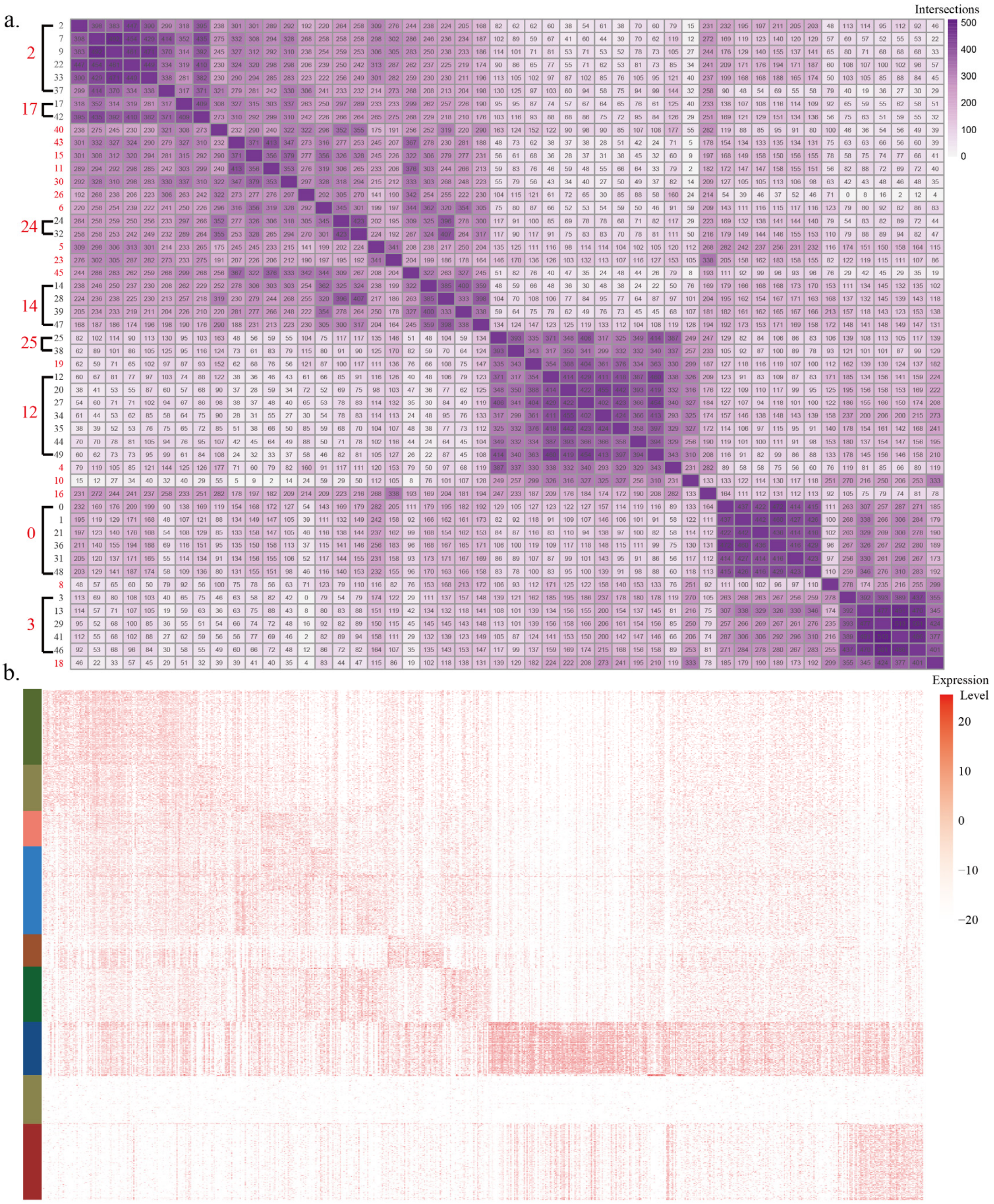


**Figure S6** Intersection and true expression of class-specific genes

**a.** The intersection of specific genes between classes, and merged with $>1500$ as the threshold (The values shown in the image are subtracted from the minimum value of all entries of the matrix, making the comparison clearer), a total of 24 final merged classes (red indexes) were obtained; **b.** Heatmap of the true expression of the specific gene in the β matrix.

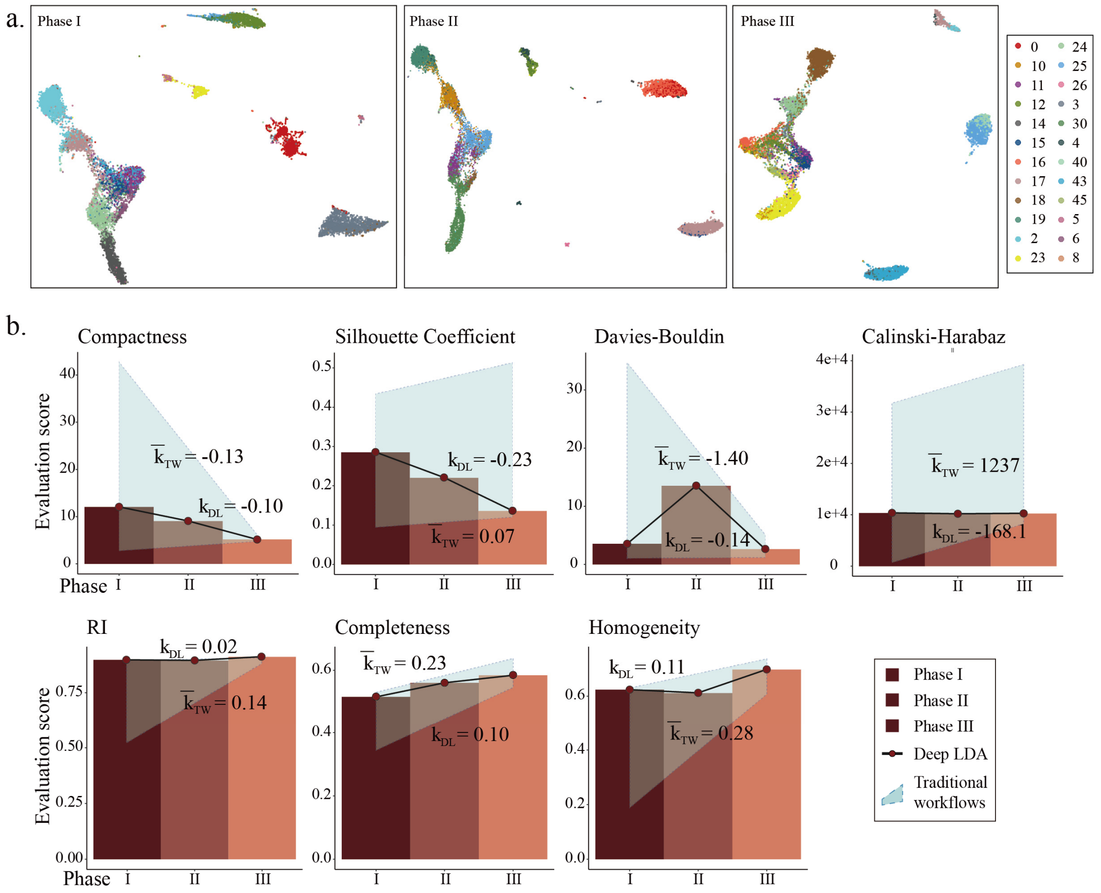


**Figure S7** Clustering results and evaluation based on Deep-LDA model

**a.** The clustering results of Deep-LDA on the three-stage data, and from left to right, 24, 19, and 21 clusters were obtained respectively; **b.** The superposition of the bar and the line plot of the Deep-LDA clustering evaluation score, and other marks are described as the same as **Fig. 5c**.

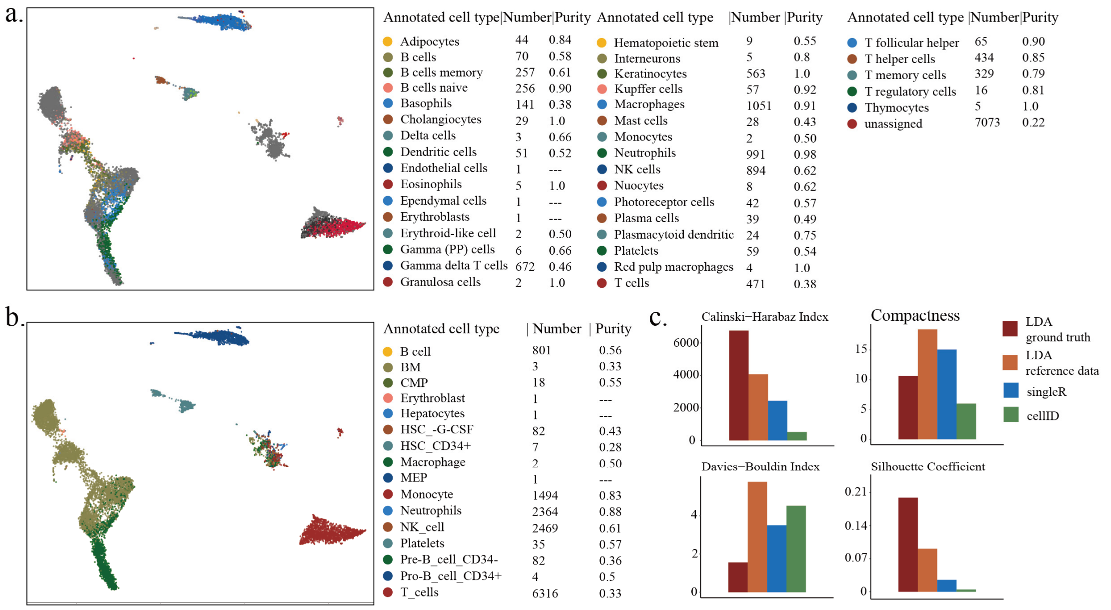


**Figure S8** Annotation results and evaluation of cellID and singleR

**a.** cellID annotation results, **b.** singleR annotation results, and marked the number and purity for each annotated cell type behind; **c.** Class evaluation score of the two methods compared with Deep-LDA

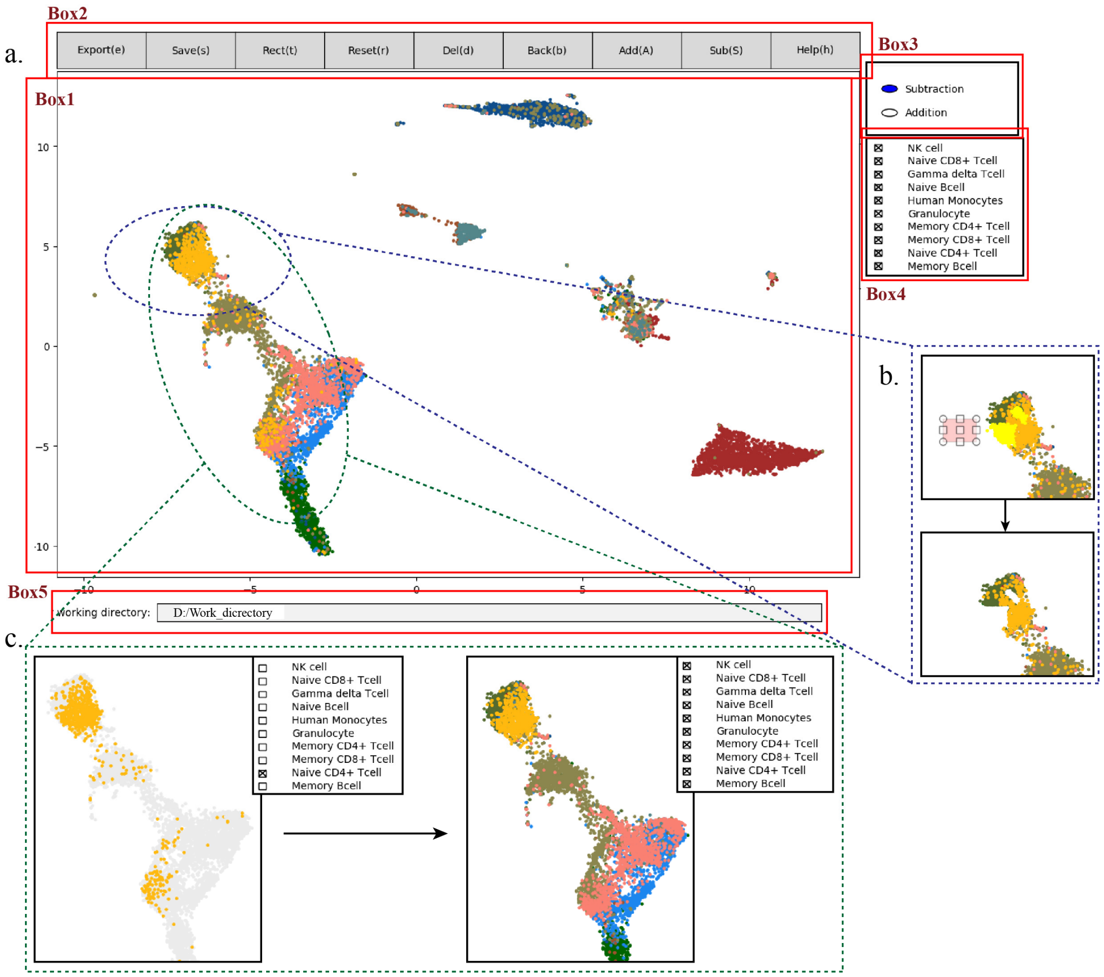


**Figure S9** Schematic diagram of the elements of the interactive cell distribution plotting tool

**a.** The interface of the interactive drawing tool. Box1 is the display module, showing the remaining cells after the operation; Box2 includes the tools that execute edit operations; Box3 is the selection mode switch operation; Box4 is the labeling activate operation; Box5 is the working directory edit panel; **b.** Shows the operation of deleting cells after pointing and boxing; **c.** Shows the operation of deleting some cells in single type of cell displaying situation (Naïve CD4+ T cells selected).
