## Supplementary Table S8 for "Optimization and redevelopment of single-cell data analysis workflow based on deep generative models"

**Supplementaty Table 8**

| Gene Information | | | | The gene expressed in scRNA-seq data | | | | The gene expressed in bulk-seq data | | | |
|---|---|---|---|---|---|---|---|---|---|---|---|
| Gene symbol | cell_type | source | Ensemble ID | count mean single.cell | logFC (count) single.cell | statistic single.cell | p.value single.cell | count mean bulk.seq | logFC (count) bulk.seq | statistic bulk.seq | p.value bulk.seq |
| PTPRC | Immune cell | Literature | ENSG00000081237.19 | NA | NA | NA | NA | NA | NA | NA | NA |
| CD3D | T cell | Literature | ENSG00000167286.9 | NA | NA | NA | NA | NA | NA | NA | NA |
| CD3E | T cell | Literature | ENSG00000198851.9 | NA | NA | NA | NA | NA | NA | NA | NA |
| CD3G | T cell | Literature | ENSG00000160654.10 | NA | NA | NA | NA | NA | NA | NA | NA |
| CD4 | CD4+ T cell | Literature | ENSG00000010610.9 | 0.423919 | 0.512036 | 29.511134 | 1.0841E-185 | 0.677972 | 1.383465 | 7.445461 | 1.20593E-07 |
| IL7R | CD4+ T cell | CellMarker | ENSG00000168685.14 | 0.594392 | 0.661508 | 44.446886 | 0 | 0.864845 | 0.502039 | 2.517078 | 0.01903377 |
| CD8A | CD8+ T cell | Literature | ENSG00000153563.15 | 0.415216 | 1.616166 | 114.355607 | 0 | 0.799061 | 0.891192 | 4.374948 | 0.00021 |
| CD8B | CD8+ T cell | Literature | ENSG00000172116.21 | 0.321179 | 1.491679 | 92.549137 | 0 | 0.767146 | 1.065816 | 5.444914 | 0.000014 |
| ABLIM1 | Naive T cell | CellMarker | ENSG00000099204.19 | 0.432929138 | 0.871387577 | 42.03304441 | 0 | 0.976329783 | 0.143304661 | 2.564221499 | 0.017142066 |
| ACTN1 | Naive T cell | CellMarker | ENSG00000072110.13 | 0.411372824 | 0.275892595 | 12.46557657 | 1.7831E-35 | 0.884091278 | 0.511201393 | 2.872078864 | 0.008474106 |
| C1orf162 | Naive T cell | CellMarker | ENSG00000143110.11 | 0.383682214 | -0.192487467 | -8.560167518 | 1.2446E-17 | 0.974326541 | 0.099422562 | 1.489278884 | 0.149648097 |
| C6orf48 | Naive T cell | CellMarker | ENSG00000204387.13 | NA | NA | NA | NA | 0.964268641 | 0.294636008 | 3.936270411 | 0.000633648 |
| CD55 | Naive T cell | CellMarker | ENSG00000196352.15 | 0.458056384 | 0.048512152 | 2.23474099 | 0.025450246 | 0.976229762 | 0.104182615 | 1.741865324 | 0.094543617 |
| EIF3L | Naive T cell | CellMarker | ENSG00000100129.17 | 0.219779092 | 0.173477622 | 7.30187773 | 2.99255E-13 | 0.975692536 | 0.127420645 | 2.12773895 | 0.044003915 |
| FAM117B | Naive T cell | CellMarker | ENSG00000138439.11 | 0.223958228 | 0.459893725 | 19.60554694 | 1.92899E-84 | 0.967456421 | 0.223406369 | 2.939920019 | 0.007229294 |
| GPR183 | Naive T cell | CellMarker | ENSG00000169508.6 | 0.428451601 | 0.163672484 | 7.432371842 | 1.12915E-13 | 0.966892051 | 0.144510356 | 1.697163106 | 0.102807839 |
| LDLRAP1 | Naive T cell | CellMarker | ENSG00000157978.11 | 0.29834129 | 0.719527162 | 32.01115599 | 5.5748E-217 | 0.898621781 | 0.454784731 | 2.711175393 | 0.01229013 |
| LRRN3 | Naive T cell | CellMarker | ENSG00000173114.12 | 0.137085042 | 0.809313902 | 34.93404423 | 2.5525E-256 | 0.615202997 | 1.56270608 | 8.965165789 | 4.56988E-09 |
| MAL | Naive T cell | CellMarker | ENSG00000172005.10 | 0.31161205 | 1.080764983 | 50.75096662 | 0 | 0.78989092 | 0.898920044 | 4.278570725 | 0.00026784 |
| MYC | Naive T cell | CellMarker | ENSG00000136997.17 | 0.19496056 | 0.232035157 | 9.729196706 | 2.66829E-22 | 0.87186082 | 0.445219522 | 2.248625678 | 0.034147433 |
| NELL2 | Naive T cell | CellMarker | ENSG00000184613.10 | 0.231562689 | 1.08657309 | 49.68748603 | 0 | 0.776707732 | 0.935744415 | 4.365562546 | 0.000215049 |
| NOSIP | Naive T cell | CellMarker | ENSG00000142546.13 | 0.25525788 | 0.654069353 | 28.52462609 | 5.9149E-174 | 0.971734871 | 0.219101207 | 3.472294441 | 0.002006176 |
| SERINC5 | Naive T cell | CellMarker | ENSG00000164300.16 | 0.274915138 | 0.45051272 | 19.46137612 | 3.01076E-83 | 0.941546533 | 0.403468213 | 3.658070746 | 0.001268229 |
| TCF7 | Naive T cell | CellMarker | ENSG00000081059.19 | 0.516911114 | 1.291843489 | 72.54886758 | 0 | 0.939763425 | 0.326967622 | 2.644352 | 0.014309912 |
| TMEM204 | Naive T cell | CellMarker | ENSG00000131634.13 | 0.127387013 | 0.401404328 | 16.75599904 | 2.10582E-62 | 0.662830834 | 1.189693427 | 4.852452988 | 6.2932E-05 |
| TRABD2A | Naive T cell | CellMarker | ENSG00000186854.10 | 0.30985569 | 1.224393379 | 59.02973354 | 0 | 0.858752947 | 0.737707678 | 4.282416419 | 0.000265254 |
| CCR7 | Naive T cell | Literature | ENSG00000126353.3 | 0.379595802 | 1.571474347 | 86.27058605 | 0 | 0.899508217 | 0.563135863 | 3.704511836 | 0.001130109 |
| SELL | Naive T cell | Literature | ENSG00000188404.8 | 0.644708801 | 0.22886636 | 12.29842202 | 1.39933E-34 | 0.973470282 | 0.170618598 | 2.691230155 | 0.01286301 |
| ADAM28 | B cell | Panlaodb | ENSG00000042980.12 | 0.167479008 | 1.241209986 | 39.65947781 | 0 | 0.783125665 | 1.03381818 | 5.499182174 | 1.25589E-05 |
| BANK1 | B cell | Panlaodb | ENSG00000153064.11 | 0.203803746 | 2.481386425 | 98.75654342 | 0 | 0.75671931 | 1.180906645 | 6.737435905 | 6.29357E-07 |
| BLK | B cell | Panlaodb | ENSG00000136573.13 | 0.10549723 | 1.163933568 | 36.59367638 | 6.0701E-280 | 0.732320678 | 1.216211574 | 6.468474959 | 1.18862E-06 |
| CD180 | B cell | Panlaodb | ENSG00000134061.5 | 0.104045258 | 0.573852123 | 17.41326821 | 3.39326E-67 | 0.51854573 | 1.527814671 | 6.258986488 | 1.96198E-06 |
| CD19 | B cell | CellMarker | ENSG00000177455.12 | 0.108221832 | 1.350190493 | 43.19107653 | 0 | 0.593934285 | 1.421791767 | 6.125537555 | 2.70684E-06 |
| CD1C | B cell | Panlaodb | ENSG00000158481.12 | 0.125047786 | 0.633271552 | 19.3100668 | 5.26578E-82 | 0.561031382 | 1.006842362 | 3.103723859 | 0.004910867 |
| CD22 | B cell | Panlaodb | ENSG00000012124.16 | 0.184483223 | 2.369420076 | 90.93955623 | 0 | 0.635022404 | 1.605435093 | 11.69048925 | 2.71296E-11 |
| CD37 | B cell | Panlaodb | ENSG00000104894.11 | 0.625461151 | 0.932398653 | 37.30539569 | 2.3938E-290 | 0.975724656 | 0.146747459 | 2.627759329 | 0.014871975 |
| CD38 | B cell | CellMarker | ENSG00000004468.12 | 0.194382147 | 0.049310892 | 1.500705812 | 0.133454454 | 0.842455034 | 0.148047394 | 0.615072713 | 0.544400488 |
| CD72 | B cell | Panlaodb | ENSG00000137101.12 | 0.127340552 | 1.015162619 | 31.64003653 | 3.3564E-212 | 0.77132368 | 0.348337052 | 1.233917893 | 0.229397344 |
| CD79A | B cell | Panlaodb | ENSG00000105369.9 | 0.25090793 | 2.862455417 | 133.3775995 | 0 | 0.791428583 | 1.054910059 | 6.038980061 | 3.33862E-06 |
| CD79B | B cell | Panlaodb | ENSG00000007312.12 | 0.126664715 | 1.390188832 | 44.76286723 | 0 | 0.908247114 | 0.625764416 | 4.918665416 | 5.35543E-05 |
| CR2 | B cell | Panlaodb | ENSG00000117322.17 | 0.081893923 | 0.549580446 | 16.626758 | 1.75662E-61 | 0.660816983 | 1.254498668 | 5.404475374 | 1.58871E-05 |
| CXCR5 | B cell | Panlaodb | ENSG00000160683.4 | 0.012077594 | -0.012984929 | -0.387715974 | 0.698232189 | 0.514086689 | 1.604414538 | 7.143358839 | 2.45188E-07 |
| FCRL2 | B cell | Panlaodb | ENSG00000132704.15 | 0.133615749 | 1.651119983 | 54.87775218 | 0 | 0.581456024 | 1.691291164 | 11.50901581 | 3.72073E-11 |
| FRK | B cell | Panlaodb | ENSG00000111816.7 | 0.008540149 | -0.009181731 | -0.274145954 | 0.783976547 | 0.510708698 | -0.42891474 | -1.097811062 | 0.283376862 |
| GPR18 | B cell | Panlaodb | ENSG00000125245.12 | 0.184786505 | 0.475246575 | 14.54507031 | 1.40662E-47 | 0.936660254 | 0.117710431 | 0.825143915 | 0.417564036 |
| GUSBP11 | B cell | Panlaodb | ENSG00000228315.11 | NA | NA | NA | NA | 0.802341768 | -0.627068284 | -2.623916355 | 0.015001678 |
| HHEX | B cell | Panlaodb | ENSG00000152804.10 | 0.203212913 | 0.660959873 | 20.45200429 | 1.29194E-91 | 0.827970078 | 0.824444123 | 4.375632285 | 0.000210493 |
| HLA-DOB | B cell | Panlaodb | ENSG00000241106.7 | 0.045990108 | 0.487364386 | 14.67912059 | 2.02545E-48 | 0.637799154 | 1.455978104 | 7.521306185 | 1.03897E-07 |
| IGHD | B cell | Panlaodb | ENSG00000211898.7 | 0.142283166 | 1.85286516 | 63.49413059 | 0 | 0.582477177 | 1.6380179 | 9.643405845 | 1.18885E-09 |
| IGHM | B cell | Panlaodb | ENSG00000211899.10 | 0.170274727 | 2.228371094 | 82.40484714 | 0 | 0.794823556 | 1.103627417 | 7.003283227 | 3.38648E-07 |
| IGKC | B cell | Panlaodb | ENSG00000211592.8 | 0.247925333 | 1.979837483 | 71.31704723 | 0 | 0.846309403 | 0.906293107 | 6.036343933 | 3.36006E-06 |
| IGLL3P | B cell | Panlaodb | ENSG00000206066.3 | NA | NA | NA | NA | NA | NA | NA | NA |
| IRF8 | B cell | Panlaodb | ENSG00000140968.10 | 0.295360578 | 0.899613122 | 28.93655051 | 8.165E-179 | 0.895536961 | 0.608256574 | 4.074062031 | 0.000449836 |
| LTB | B cell | Panlaodb | ENSG00000227507.2 | 0.069207118 | 0.117498955 | 3.518035253 | 0.000436147 | 0.958521128 | 0.21038256 | 2.137806825 | 0.043116982 |
| LY86 | B cell | Panlaodb | ENSG00000112799.8 | 0.252000714 | 0.390245366 | 12.09922316 | 1.5734E-33 | 0.764393873 | 1.047049239 | 5.187881037 | 2.72638E-05 |
| MS4A1 | B cell | Panlaodb | ENSG00000156738.17 | 0.235632401 | 2.844756876 | 130.3715243 | 0 | 0.802066382 | 1.112132526 | 7.599409694 | 8.72086E-08 |
| NMBR | B cell | Panlaodb | ENSG00000135577.4 | NA | NA | NA | NA | NA | NA | NA | NA |
| P2RX5 | B cell | Panlaodb | ENSG00000083454.21 | 0.198497539 | 1.66805682 | 56.31799027 | 0 | 0.909973234 | 0.666795062 | 5.798213305 | 6.0081E-06 |
| PNOC | B cell | Panlaodb | ENSG00000168081.8 | 0.108714764 | 1.077686969 | 33.66342537 | 7.2425E-239 | 0.582565553 | 1.688553863 | 11.46380096 | 4.02762E-11 |
| PTPRCAP | B cell | Panlaodb | ENSG00000213402.2 | NA | NA | NA | NA | NA | NA | NA | NA |
| RALGPS2 | B cell | Panlaodb | ENSG00000116191.17 | 0.237186956 | 2.314907618 | 89.25907664 | 0 | 0.830031875 | 0.864053474 | 4.838555399 | 6.55115E-05 |
| SIK1 | B cell | Panlaodb | ENSG00000142178.8 | 0.055346489 | -0.039228906 | -1.173079815 | 0.240783965 | 0.947093789 | -0.090191809 | -0.709812066 | 0.484785012 |
| SLC12A1 | B cell | Panlaodb | ENSG00000074803.18 | NA | NA | NA | NA | 0.337246647 | 0.052058194 | 0.118307068 | 0.906826931 |
| SPIB | B cell | Panlaodb | ENSG00000269404.6 | 0.140025326 | 1.674192939 | 55.87609391 | 0 | 0.575175938 | 1.674581388 | 10.46091484 | 2.47742E-10 |
| STAP1 | B cell | Panlaodb | ENSG00000035720.7 | 0.114216936 | 0.955572559 | 29.61074982 | 6.7683E-187 | 0.789754199 | 0.732030224 | 3.104651915 | 0.004899997 |
| VPREB3 | B cell | Panlaodb | ENSG00000128218.7 | 0.125981141 | 1.708663129 | 57.16254474 | 0 | 0.590754301 | 1.606676946 | 9.129830801 | 3.32289E-09 |
| EEF1B2 | Naive CD4+ T cell | CellMarker | ENSG00000114942.13 | 0.697830571 | 0.391253378 | 14.43814425 | 6.51911E-47 | 0.97706834 | 0.046361271 | 0.578624027 | 0.56831535 |
| FHIT | Naive CD4+ T cell | CellMarker | ENSG00000189283.9 | 0.135351485 | 0.936314796 | 25.4876181 | 4.4522E-140 | 0.854486918 | 0.57311555 | 2.033284539 | 0.053398698 |
| GIMAP5 | Naive CD4+ T cell | CellMarker | ENSG00000196329.11 | 0.0926435 | 0.285876022 | 7.584298764 | 3.55524E-14 | 0.568703678 | 1.625456545 | 4.361673772 | 0.00021676 |
| GIMAP8 | Naive CD4+ T cell | CellMarker | ENSG00000171115.3 | 0.205909526 | 0.156050718 | 4.205693121 | 2.61919E-05 | 0.723779348 | 0.805705114 | 2.111031471 | 0.045540223 |
| PRKCA | Naive CD4+ T cell | CellMarker | ENSG00000154229.11 | 0.176930013 | 0.358823683 | 9.641687161 | 6.24623E-22 | 0.801198808 | 0.737180778 | 2.280663653 | 0.031885826 |
| RPS5 | Naive CD4+ T cell | CellMarker | ENSG00000083845.8 | 0.786518829 | 0.485290871 | 20.86612496 | 3.14949E-95 | 0.977726945 | 0.053641065 | 0.698879053 | 0.491446905 |
| RSL1D1 | Naive CD4+ T cell | CellMarker | ENSG00000171490.12 | 0.506857289 | 0.303197564 | 9.284694973 | 1.85908E-20 | 0.977393827 | 0.089319654 | 1.160176736 | 0.257560227 |
| SATB1 | Naive CD4+ T cell | CellMarker | ENSG00000182568.16 | 0.518001158 | 0.644633716 | 20.11759863 | 9.52996E-89 | 0.959681235 | 0.236473649 | 1.804720636 | 0.08387402 |
| SLC40A1 | Naive CD4+ T cell | CellMarker | ENSG00000138449.10 | 0.181584359 | 0.031937849 | 0.856066762 | 0.391975654 | 0.873206199 | 0.6217358 | 2.448399765 | 0.02216124 |
| SVIL | Naive CD4+ T cell | CellMarker | ENSG00000197321.14 | 0.204581268 | 0.100232457 | 2.699573022 | 0.006951276 | 0.876727464 | 0.401962299 | 1.502023559 | 0.146332109 |
| TESPA1 | Naive CD4+ T cell | CellMarker | ENSG00000135426.15 | 0.292331278 | 0.546710688 | 15.18863356 | 1.09611E-51 | 0.907608842 | 0.442575247 | 2.002318901 | 0.056850409 |
| TSHZ2 | Naive CD4+ T cell | CellMarker | ENSG00000182463.15 | 0.116847825 | 0.792929654 | 21.39090207 | 6.6495E-100 | 0.642097494 | 1.384581396 | 3.76580652 | 0.000968985 |
| MAF | Memory CD4+ T cell | SC2Disease | ENSG00000178573.6 | 0.388455383 | 0.730930059 | 38.80345896 | 8.85268861804e-313 | 0.781714909 | 0.735992465 | 2.156758175 | 0.041415867 |
| IGHG4 | Memory CD4+ T cell | SC2Disease | ENSG00000211892.4 | 0.029847098 | -0.028282843 | -1.315877582 | 0.188236901 | 0.345727494 | -0.395117136 | -0.695406404 | 0.493579203 |
| STAT1 | Memory CD4+ T cell | SC2Disease | ENSG00000115415.18 | 0.403153806 | 0.006353038 | 0.322507473 | 0.747073143 | 0.975000427 | -0.019496031 | -0.217313029 | 0.829828727 |
| XAF1 | Memory CD4+ T cell | SC2Disease | ENSG00000132530.16 | 0.119027423 | -0.028631441 | -1.340946565 | 0.17995978 | 0.938557891 | -0.115128056 | -0.620105945 | 0.541118954 |
| IFI44L | Memory CD4+ T cell | SC2Disease | ENSG00000137959.15 | 0.146674916 | 0.022607329 | 1.062708538 | 0.287932666 | 0.797823125 | 0.595117384 | 1.758399126 | 0.091618498 |
| EPSTI1 | Memory CD4+ T cell | SC2Disease | ENSG00000133106.14 | 0.219109884 | 0.031127611 | 1.483286285 | 0.138021106 | 0.886458287 | 0.243288645 | 0.92422459 | 0.3647061 |
| IFI27 | Memory CD4+ T cell | SC2Disease | ENSG00000165949.12 | 0.014791971 | 0.044144719 | 2.053362363 | 0.040056174 | 0.199826313 | 1.598610508 | 3.183691486 | 0.004043856 |
| AIF1 | Naive CD8+ T cell | CellMarker | ENSG00000204472.12 | 0.397702369 | 0.041598376 | 1.513869345 | 0.130081798 | 0.819511871 | 0.402074537 | 1.217232775 | 0.235526307 |
| APBA2 | Naive CD8+ T cell | CellMarker | ENSG00000034053.14 | 0.090245328 | 0.294655164 | 9.921960449 | 3.99076E-23 | 0.833835721 | 0.553821158 | 1.808125579 | 0.083325758 |
| APEX1 | Naive CD8+ T cell | CellMarker | ENSG00000100823.11 | 0.374544726 | 0.297121 | 10.74483251 | 7.99469E-27 | 0.972379721 | 0.080550123 | 0.815915826 | 0.422692747 |
| ARHGAP45 | Naive CD8+ T cell | CellMarker | ENSG00000180448.10 | 0.292693269 | 0.138577366 | 4.845034812 | 1.2796E-06 | 0.975123301 | -0.00729555 | -0.081810969 | 0.935485687 |
| ARHGEF18 | Naive CD8+ T cell | CellMarker | ENSG00000104880.17 | 0.032754423 | 0.067270539 | 2.249687621 | 0.024484418 | 0.80318967 | 0.256141288 | 0.729113182 | 0.473093969 |
| ATM | Naive CD8+ T cell | CellMarker | ENSG00000149311.18 | 0.573063071 | 0.640458458 | 26.71782962 | 2.182E-153 | 0.976700297 | 0.086980046 | 1.083508713 | 0.289509129 |
| BEX4 | Naive CD8+ T cell | CellMarker | ENSG00000102409.9 | 0.284800359 | 0.291514122 | 10.19652229 | 2.50462E-24 | 0.954951973 | 0.09899387 | 0.658565331 | 0.516537527 |
| CA6 | Naive CD8+ T cell | CellMarker | ENSG00000131686.14 | 0.086708444 | 0.750194715 | 25.75999597 | 5.647E-143 | 0.460022067 | 2.184523987 | 7.202851329 | 2.09919E-07 |
| CAMK4 | Naive CD8+ T cell | CellMarker | ENSG00000152495.10 | 0.346747804 | 0.765447978 | 28.01551675 | 4.8719E-168 | 0.853935648 | 0.440573766 | 1.507069725 | 0.145038999 |
| CD248 | Naive CD8+ T cell | CellMarker | ENSG00000174807.3 | 0.091060584 | 0.864615418 | 29.93752855 | 7.1964E-191 | 0.4628338 | 1.792070579 | 4.522964075 | 0.000144179 |
| CEP68 | Naive CD8+ T cell | CellMarker | ENSG00000011523.13 | 0.194805775 | 0.297672848 | 10.17798486 | 3.02633E-24 | 0.888761282 | 0.374744481 | 1.478281707 | 0.152532092 |
| COQ8A | Naive CD8+ T cell | CellMarker | ENSG00000163050.16 | 0.218079429 | 0.075394524 | 2.581703901 | 0.009841566 | 0.959010392 | 0.049264322 | 0.349195824 | 0.730033245 |
| DGKA | Naive CD8+ T cell | CellMarker | ENSG00000065357.19 | 0.453131274 | 0.720524636 | 27.71501493 | 1.3667E-164 | 0.970752177 | 0.167690359 | 1.672362807 | 0.107636381 |
| EEF2 | Naive CD8+ T cell | CellMarker | ENSG00000167658.15 | 0.732805492 | 0.418715489 | 20.76481847 | 2.4492E-94 | 0.975308417 | 0.114885608 | 1.348641508 | 0.190229681 |
| EIF3E | Naive CD8+ T cell | CellMarker | ENSG00000104408.9 | 0.606642989 | 0.641218044 | 27.60838034 | 2.245E-163 | 0.977193235 | 0.102849788 | 1.333979304 | 0.194918944 |
| EPHX2 | Naive CD8+ T cell | CellMarker | ENSG00000120915.13 | 0.122539683 | 0.51836729 | 17.64846272 | 5.9356E-69 | 0.761060172 | 0.871930437 | 2.522109901 | 0.018823809 |
| EXOSC8 | Naive CD8+ T cell | CellMarker | ENSG00000120699.12 | 0.196522936 | 0.163169852 | 5.566470095 | 2.64781E-08 | 0.962750543 | 0.081832116 | 0.628673129 | 0.535590078 |
| FAM102A | Naive CD8+ T cell | CellMarker | ENSG00000167106.11 | 0.233967575 | 0.569088875 | 19.83119221 | 2.51577E-86 | 0.903111583 | 0.323327345 | 1.368304016 | 0.184080595 |
| FKBP5 | Naive CD8+ T cell | CellMarker | ENSG00000096060.14 | 0.341871679 | 0.455841195 | 16.35547996 | 1.4367E-59 | 0.943922277 | 0.179701197 | 1.041975558 | 0.307961191 |
| GCSAM | Naive CD8+ T cell | CellMarker | ENSG00000174500.12 | 0.017080297 | 0.033682735 | 1.125841151 | 0.26025227 | 0.664525975 | 0.88199384 | 2.126740195 | 0.044082852 |
| GIMAP2 | Naive CD8+ T cell | CellMarker | ENSG00000106560.10 | 0.292384557 | 0.287004752 | 10.06155043 | 9.85715E-24 | 0.899197469 | 0.305743765 | 1.258380326 | 0.220539549 |
| GPR155 | Naive CD8+ T cell | CellMarker | ENSG00000163328.13 | 0.229151705 | 0.331657529 | 11.43703462 | 3.72888E-30 | 0.897040886 | 0.420883963 | 1.763014544 | 0.090820913 |
| IGF1R | Naive CD8+ T cell | CellMarker | ENSG00000140443.14 | 0.209825889 | 0.138597908 | 4.739966751 | 2.1588E-06 | 0.877188565 | 0.50557436 | 1.948932781 | 0.063265073 |
| ITFG2 | Naive CD8+ T cell | CellMarker | ENSG00000111203.12 | 0.225410205 | 0.061997547 | 2.126387469 | 0.033488642 | 0.953823347 | 0.085696404 | 0.558638028 | 0.581661621 |
| LDHB | Naive CD8+ T cell | CellMarker | ENSG00000111716.12 | 0.260802262 | 0.517714318 | 18.12632728 | 1.36078E-72 | 0.965504627 | 0.244283632 | 2.172039732 | 0.040115822 |
| LEF1 | Naive CD8+ T cell | CellMarker | ENSG00000138795.9 | 0.387785024 | 1.591885447 | 66.14949429 | 0 | 0.83854936 | 0.642280447 | 2.186430337 | 0.038924719 |
| LMO7 | Naive CD8+ T cell | CellMarker | ENSG00000136153.19 | 0.110046605 | 0.336335242 | 11.36039136 | 8.919E-30 | 0.87888586 | 0.51305291 | 1.999796814 | 0.057138468 |
| LRRC75A | Naive CD8+ T cell | CellMarker | ENSG00000181350.11 | 0.0560015 | -0.04550699 | -1.523270603 | 0.127713849 | 0.364274906 | -0.416314178 | -0.739404121 | 0.46694007 |
| MAML2 | Naive CD8+ T cell | CellMarker | ENSG00000184384.13 | 0.262286933 | 0.47848405 | 16.73092901 | 3.18115E-62 | 0.909204338 | 0.347815364 | 1.542095325 | 0.136329967 |
| MLXIP | Naive CD8+ T cell | CellMarker | ENSG00000175727.13 | 0.249073556 | 0.406915452 | 14.13739767 | 4.59548E-45 | 0.931919695 | 0.231957009 | 1.197183478 | 0.243101155 |
| NDFIP1 | Naive CD8+ T cell | CellMarker | ENSG00000131507.10 | 0.449785848 | 0.533444803 | 20.22755153 | 1.10061E-89 | 0.97405051 | 0.15770477 | 1.795901562 | 0.085302235 |
| NOG | Naive CD8+ T cell | CellMarker | ENSG00000183691.4 | 0.089265755 | 0.535401164 | 18.17630007 | 5.59555E-73 | 0.514004791 | 1.505796133 | 3.573210309 | 0.001562686 |
| NPAT | Naive CD8+ T cell | CellMarker | ENSG00000149308.16 | 0.315676684 | 0.465688276 | 16.55376511 | 5.78713E-61 | 0.975361437 | 0.143763083 | 1.730606819 | 0.096555267 |
| NPM1 | Naive CD8+ T cell | CellMarker | ENSG00000181163.13 | 0.717262383 | 0.810924893 | 40.96950383 | 0 | 0.977664377 | 0.081533856 | 1.074817754 | 0.293303319 |
| OXNAD1 | Naive CD8+ T cell | CellMarker | ENSG00000154814.13 | 0.302017346 | 0.990682812 | 36.32799808 | 4.2656E-276 | 0.946605073 | 0.323690642 | 2.058924216 | 0.050681392 |
| PASK | Naive CD8+ T cell | CellMarker | ENSG00000115687.13 | 0.207629349 | 0.775753977 | 27.19443443 | 1.0719E-158 | 0.916592829 | 0.464309883 | 2.275006209 | 0.032272024 |
| PDE3B | Naive CD8+ T cell | CellMarker | ENSG00000152270.8 | 0.373076354 | 0.898401419 | 33.62814662 | 2.1917E-238 | 0.911725652 | 0.345973213 | 1.56055763 | 0.131915031 |
| PDK1 | Naive CD8+ T cell | CellMarker | ENSG00000152256.13 | 0.204993858 | 0.400533793 | 13.76595942 | 7.81659E-43 | 0.958372309 | 0.27625142 | 2.103260252 | 0.046274783 |
| PIK3IP1 | Naive CD8+ T cell | CellMarker | ENSG00000100100.12 | 0.519564999 | 0.872894069 | 35.60458227 | 9.3846E-266 | 0.972806131 | 0.129009391 | 1.360175158 | 0.186603498 |
| PLEKHB1 | Naive CD8+ T cell | CellMarker | ENSG00000021300.13 | 0.175941516 | 0.393468251 | 13.44224503 | 6.17234E-41 | 0.808847178 | 0.589778626 | 1.795043127 | 0.085442537 |
| PSIP1 | Naive CD8+ T cell | CellMarker | ENSG00000164985.14 | 0.459122217 | 0.3218498 | 12.15580058 | 7.94468E-34 | 0.977609062 | 0.086632072 | 1.14147332 | 0.265102893 |
| R3HDM4 | Naive CD8+ T cell | CellMarker | ENSG00000198858.9 | 0.521348972 | -0.062966783 | -2.461951554 | 0.013830424 | 0.976361136 | 0.030106356 | 0.360301029 | 0.721820917 |
| RACK1 | Naive CD8+ T cell | CellMarker | ENSG00000204628.11 | 0.791049654 | 0.534433326 | 29.87792267 | 3.8474E-190 | 0.978121035 | 0.059628119 | 0.801125007 | 0.431037136 |
| RCAN3 | Naive CD8+ T cell | CellMarker | ENSG00000117602.11 | 0.423212848 | 1.246040316 | 49.86119596 | 0 | 0.938934059 | 0.332654164 | 1.918644568 | 0.067181648 |
| RIC3 | Naive CD8+ T cell | CellMarker | ENSG00000166405.14 | 0.127695611 | 0.213823247 | 7.217588834 | 5.56707E-13 | 0.892401488 | 0.372820065 | 1.499510532 | 0.146976911 |
| RPS8 | Naive CD8+ T cell | CellMarker | ENSG00000142937.11 | 0.83960278 | 0.558761646 | 35.49341022 | 3.5197E-264 | 0.977551178 | 0.068807631 | 0.894209419 | 0.38022281 |
| SCML4 | Naive CD8+ T cell | CellMarker | ENSG00000146285.13 | 0.286802686 | 0.504995633 | 17.8090302 | 3.63737E-70 | 0.97006776 | 0.142357888 | 1.363525386 | 0.185560433 |
| SGSM3 | Naive CD8+ T cell | CellMarker | ENSG00000100359.20 | 0.034160595 | 0.119805884 | 4.008373031 | 6.14587E-05 | 0.787609122 | 0.416846602 | 1.163189245 | 0.256358514 |
| SH3YL1 | Naive CD8+ T cell | CellMarker | ENSG00000035115.21 | 0.20894781 | 0.419546338 | 14.44139844 | 6.22522E-47 | 0.954821521 | 0.176452571 | 1.195381881 | 0.243790648 |
| SMAP2 | Naive CD8+ T cell | CellMarker | ENSG00000084070.11 | 0.683039645 | 0.014190758 | 0.646550656 | 0.517933516 | 0.977732541 | 0.012040914 | 0.155733554 | 0.877565077 |
| SNED1 | Naive CD8+ T cell | CellMarker | ENSG00000162804.13 | 0.079683322 | 0.274138011 | 9.218372071 | 3.44504E-20 | 0.779675193 | 0.453342128 | 1.248298696 | 0.224142501 |
| SNURF | Naive CD8+ T cell | CellMarker | ENSG00000273173.5 | 0.062173221 | -0.010653027 | -0.356693298 | 0.721326857 | 0.779774352 | 0.105146593 | 0.280959719 | 0.781185399 |
| SPINT2 | Naive CD8+ T cell | CellMarker | ENSG00000167642.12 | 0.292230883 | 0.23703667 | 8.299772606 | 1.13887E-16 | 0.871251499 | 0.334545515 | 1.202611632 | 0.241032556 |
| SPTBN1 | Naive CD8+ T cell | CellMarker | ENSG00000115306.15 | 0.375132912 | 0.275035556 | 9.942764215 | 3.24389E-23 | 0.971340929 | 0.036492778 | 0.352045372 | 0.727922804 |
| STMN3 | Naive CD8+ T cell | CellMarker | ENSG00000197457.9 | 0.187636307 | 0.467195283 | 16.03937901 | 2.22781E-57 | 0.831893291 | 0.561644283 | 1.825106436 | 0.080645741 |
| THAP7 | Naive CD8+ T cell | CellMarker | ENSG00000184436.11 | 0.144212549 | 0.135035603 | 4.563313717 | 5.07852E-06 | 0.918827939 | 0.202592552 | 0.930054536 | 0.36174247 |
| THEM4 | Naive CD8+ T cell | CellMarker | ENSG00000159445.12 | 0.274666574 | 0.363702982 | 12.70900335 | 8.45793E-37 | 0.974919402 | 0.139244601 | 1.6310493 | 0.116130728 |
| TMEM123 | Naive CD8+ T cell | CellMarker | ENSG00000152558.14 | 0.611739621 | 0.614996796 | 26.55649691 | 1.3085E-151 | 0.976341955 | 0.136026428 | 1.72029665 | 0.098443071 |
| TMEM63A | Naive CD8+ T cell | CellMarker | ENSG00000196187.11 | 0.23915324 | 0.25258195 | 8.714391586 | 3.25153E-18 | 0.951300797 | 0.129030434 | 0.816106723 | 0.422585712 |
| TOMM7 | Naive CD8+ T cell | CellMarker | ENSG00000196683.10 | 0.696487233 | 0.489221684 | 23.11308125 | 5.2903E-116 | 0.975831388 | 0.104036232 | 1.246810189 | 0.224678234 |
| TPCN1 | Naive CD8+ T cell | CellMarker | ENSG00000186815.12 | 0.197639997 | 0.04441866 | 1.514100042 | 0.130023286 | 0.831901317 | 0.321504454 | 0.998556841 | 0.328120287 |
| TRAP1 | Naive CD8+ T cell | CellMarker | ENSG00000126602.10 | 0.074975941 | 0.101439019 | 3.400835117 | 0.000673701 | 0.947905549 | 0.203897832 | 1.254038233 | 0.222085863 |
| TRMT1 | Naive CD8+ T cell | CellMarker | ENSG00000104907.12 | 0.180874774 | 0.091744955 | 3.117977086 | 0.0018247 | 0.961923642 | 0.045392591 | 0.34066445 | 0.736364741 |
| UBQLN2 | Naive CD8+ T cell | CellMarker | ENSG00000188021.8 | 0.311225448 | 0.078637973 | 2.764615115 | 0.005706536 | 0.972958265 | 0.019256791 | 0.196847997 | 0.845628859 |
| VIPR1 | Naive CD8+ T cell | CellMarker | ENSG00000114812.12 | 0.190297101 | 0.463867136 | 15.93139173 | 1.22135E-56 | 0.693632923 | 1.06380671 | 2.825644248 | 0.009434857 |
| WDR43 | Naive CD8+ T cell | CellMarker | ENSG00000163811.11 | 0.343889612 | 0.186295532 | 6.636275676 | 3.33669E-11 | 0.973372976 | 0.06602157 | 0.692510721 | 0.495362851 |
| ZNF101 | Naive CD8+ T cell | CellMarker | ENSG00000181896.11 | 0.332441228 | 0.674610668 | 24.39749839 | 9.3116E-129 | 0.973181179 | 0.17848025 | 1.986741672 | 0.058658766 |
| ADAM8 | Memory CD8+ T cell | CellMarker | ENSG00000151651.15 | 0.470861219 | -0.130312428 | -5.267053932 | 1.40705E-07 | 0.938066302 | -0.027608871 | -0.146973673 | 0.884398798 |
| ADRB2 | Memory CD8+ T cell | CellMarker | ENSG00000169252.5 | 0.17477441 | 0.207645119 | 7.537523237 | 5.08653E-14 | 0.814413415 | 0.295247195 | 0.869033798 | 0.393564664 |
| AGTPBP1 | Memory CD8+ T cell | CellMarker | ENSG00000135049.15 | 0.364427836 | -0.078198352 | -2.99427405 | 0.002755844 | 0.952763628 | 0.096415564 | 0.620030328 | 0.541167798 |
| AHNAK | Memory CD8+ T cell | CellMarker | ENSG00000124942.13 | 0.718049452 | 0.32284972 | 16.62170764 | 1.9077E-61 | 0.971619118 | 0.035154658 | 0.343220488 | 0.734465521 |
| ARHGAP25 | Memory CD8+ T cell | CellMarker | ENSG00000163219.11 | 0.3999635 | 0.004124781 | 0.160391883 | 0.872574721 | 0.976704232 | -0.090613658 | -1.135769033 | 0.267434791 |
| ARL4C | Memory CD8+ T cell | CellMarker | ENSG00000188042.7 | 0.754320207 | 0.52937747 | 29.45380196 | 5.3208E-185 | 0.966815307 | 0.100875709 | 0.861411482 | 0.397663208 |
| BIN2 | Memory CD8+ T cell | CellMarker | ENSG00000110934.10 | 0.552167483 | -0.034204876 | -1.46039797 | 0.144203401 | 0.974501927 | -0.025900659 | -0.283518812 | 0.779246827 |
| C12orf75 | Memory CD8+ T cell | CellMarker | ENSG00000235162.8 | 0.396129644 | 0.37848692 | 14.80684706 | 3.14518E-49 | 0.966830322 | 0.137305109 | 1.189003901 | 0.246242355 |
| C1orf21 | Memory CD8+ T cell | CellMarker | ENSG00000116667.13 | 0.31878003 | 0.138368019 | 5.209920426 | 1.91631E-07 | 0.813844836 | 0.424011466 | 1.267737228 | 0.217234294 |
| CAPN2 | Memory CD8+ T cell | CellMarker | ENSG00000162909.17 | 0.398189252 | 0.085774052 | 3.333878672 | 0.000858685 | 0.975405243 | -0.01195495 | -0.136355749 | 0.892694379 |
| CASP8 | Memory CD8+ T cell | CellMarker | ENSG00000064012.21 | 0.485703428 | 0.064486085 | 2.628249548 | 0.008592016 | 0.977456121 | 0.019500545 | 0.249104573 | 0.805431037 |
| CD300A | Memory CD8+ T cell | CellMarker | ENSG00000167851.14 | 0.352628395 | -0.121964462 | -4.649880438 | 3.35194E-06 | 0.862117802 | 0.163710948 | 0.55486487 | 0.584198383 |
| CEP78 | Memory CD8+ T cell | CellMarker | ENSG00000148019.13 | 0.332534712 | 0.030163671 | 1.140318889 | 0.254173179 | 0.934383448 | 0.1348683 | 0.699102425 | 0.491308363 |
| DSTN | Memory CD8+ T cell | CellMarker | ENSG00000125868.15 | 0.359472637 | 0.015345898 | 0.586226425 | 0.557732914 | 0.95764068 | 0.044750176 | 0.309643878 | 0.759544198 |
| EFHD2 | Memory CD8+ T cell | CellMarker | ENSG00000142634.12 | 0.673658909 | -0.006258549 | -0.300825041 | 0.763552425 | 0.957057195 | 0.004819144 | 0.032941337 | 0.973997868 |
| EMP3 | Memory CD8+ T cell | CellMarker | ENSG00000142227.10 | 0.642118268 | -0.05528127 | -2.563912422 | 0.010360414 | 0.977012092 | 0.029873762 | 0.372200724 | 0.713058536 |
| FCRL6 | Memory CD8+ T cell | CellMarker | ENSG00000181036.13 | 0.296529479 | 0.187872918 | 7.026837275 | 2.21134E-12 | 0.831662246 | 0.285901227 | 0.883676853 | 0.385767763 |
| FGR | Memory CD8+ T cell | CellMarker | ENSG00000000938.12 | 0.541124391 | -0.377645962 | -16.135853 | 4.82387E-58 | 0.914525132 | -0.032066585 | -0.140432795 | 0.889507528 |

| | | | | | | | | | | | |
|---|---|---|---|---|---|---|---|---|---|---|---|
| FLNA | Memory CD8+ T cell | CellMarker | ENSG00000196924.15 | 0.411701983 | 0.062449862 | 2.442601554 | 0.01459423 | 0.971968049 | 0.077415513 | 0.773087913 | 0.44713045 |
| FUT11 | Memory CD8+ T cell | CellMarker | ENSG00000196968.10 | 0.210497542 | 0.018855765 | 0.687926188 | 0.491510778 | 0.887225604 | 0.180156408 | 0.681756692 | 0.50201671 |
| GAB3 | Memory CD8+ T cell | CellMarker | ENSG00000160219.11 | 0.240290242 | 0.183687462 | 6.759655658 | 1.43852E-11 | 0.957962111 | 0.083974972 | 0.587441394 | 0.562475689 |
| GK5 | Memory CD8+ T cell | CellMarker | ENSG00000175066.15 | 0.268166782 | 0.014456142 | 0.535109056 | 0.592583035 | 0.927035909 | 0.151124176 | 0.734716437 | 0.469736794 |
| GLUL | Memory CD8+ T cell | CellMarker | ENSG00000135821.18 | 0.447600229 | -0.368526453 | -14.79847534 | 3.55522E-49 | 0.915637922 | -0.070135316 | -0.309994666 | 0.759280739 |
| KCNAB2 | Memory CD8+ T cell | CellMarker | ENSG00000069424.14 | 0.439123074 | 0.00714616 | 0.283407735 | 0.776868534 | 0.952009995 | 0.002039908 | 0.012868427 | 0.989840729 |
| LGALS1 | Memory CD8+ T cell | CellMarker | ENSG00000100097.11 | 0.39665299 | -0.166329927 | -6.46745703 | 1.03009E-10 | 0.970410273 | 0.060146733 | 0.565860785 | 0.576819717 |
| LITAF | Memory CD8+ T cell | CellMarker | ENSG00000189067.12 | 0.771149342 | 0.110086426 | 6.130911082 | 8.9749E-10 | 0.961804651 | 0.051316151 | 0.384963178 | 0.703705763 |
| LLGL2 | Memory CD8+ T cell | CellMarker | ENSG00000073350.13 | 0.185364875 | 0.028757474 | 1.043838247 | 0.296578441 | 0.941871711 | 0.101175326 | 0.564526573 | 0.577712525 |
| LPCAT1 | Memory CD8+ T cell | CellMarker | ENSG00000153395.9 | 0.089178538 | 0.043156667 | 1.54580102 | 0.122175466 | 0.871953249 | 0.185505994 | 0.655307164 | 0.518595389 |
| LUZP6 | Memory CD8+ T cell | CellMarker | ENSG00000267697.1 | NA | NA | NA | NA | NA | NA | NA | NA |
| MBP | Memory CD8+ T cell | CellMarker | ENSG00000197971.14 | 0.630759059 | -0.084314436 | -3.865188697 | 0.00011151 | 0.973922916 | -0.041842299 | -0.447667128 | 0.658468019 |
| MTSS1 | Memory CD8+ T cell | CellMarker | ENSG00000170873.18 | 0.24925157 | -0.050886441 | -1.87408651 | 0.060939526 | 0.954587416 | -0.011693895 | -0.076736227 | 0.939479013 |
| MYO1F | Memory CD8+ T cell | CellMarker | ENSG00000142347.17 | 0.566742104 | -0.200166112 | -8.669485558 | 4.81764E-18 | 0.967086046 | 0.023259212 | 0.197149914 | 0.845395127 |
| MYO1G | Memory CD8+ T cell | CellMarker | ENSG00000136286.15 | 0.254816592 | 0.023984264 | 0.884533948 | 0.376423391 | 0.971091363 | -0.022539171 | -0.215705232 | 0.831067442 |
| OSBPL5 | Memory CD8+ T cell | CellMarker | ENSG00000021762.19 | 0.132166562 | -0.041129854 | -1.480235493 | 0.138833145 | 0.734284236 | 0.088714109 | 0.217241601 | 0.82988372 |
| PATL2 | Memory CD8+ T cell | CellMarker | ENSG00000229474.6 | 0.214041948 | 0.196254752 | 7.178852164 | 7.38766E-13 | 0.968688839 | 0.007890167 | 0.069906166 | 0.944856219 |
| PIK3R5 | Memory CD8+ T cell | CellMarker | ENSG00000141506.13 | 0.454174897 | 0.12376086 | 4.953432138 | 7.37753E-07 | 0.96422668 | 0.106792142 | 0.855171362 | 0.401038856 |
| PLEK | Memory CD8+ T cell | CellMarker | ENSG00000115956.9 | 0.573491073 | -0.285648334 | -12.477403 | 1.53916E-35 | 0.919891211 | 0.033680642 | 0.15320636 | 0.879535531 |
| PLEKHG3 | Memory CD8+ T cell | CellMarker | ENSG00000126822.16 | 0.269735931 | -0.091410837 | -3.386554919 | 0.000709728 | 0.771724512 | 0.231414258 | 0.611898404 | 0.546444362 |
| PRKCB | Memory CD8+ T cell | CellMarker | ENSG00000166501.13 | 0.593121692 | 0.001477523 | 0.065286515 | 0.947946821 | 0.976494731 | -0.065171919 | -0.796738266 | 0.433530739 |
| PRSS23 | Memory CD8+ T cell | CellMarker | ENSG00000150687.11 | 0.272290352 | 0.086274961 | 3.198502663 | 0.001384539 | 0.825091238 | 0.218096222 | 0.656599741 | 0.517778248 |
| PXN | Memory CD8+ T cell | CellMarker | ENSG00000089159.16 | 0.24391698 | -0.013993867 | -0.514603579 | 0.60683827 | 0.896295204 | 0.112521294 | 0.443353999 | 0.66153992 |
| RAP1B | Memory CD8+ T cell | CellMarker | ENSG00000127314.17 | 0.479419842 | 0.086743395 | 3.522324458 | 0.000429156 | 0.977362415 | -0.025241607 | -0.320876122 | 0.751123145 |
| RAP1GAP2 | Memory CD8+ T cell | CellMarker | ENSG00000132359.14 | 0.316153129 | -0.018515146 | -0.695836171 | 0.486543104 | 0.965816008 | 0.075451202 | 0.623372458 | 0.539007069 |
| RAP2B | Memory CD8+ T cell | CellMarker | ENSG00000181467.4 | 0.492971356 | -0.013335594 | -0.545899838 | 0.585143577 | 0.974286338 | -0.002668129 | -0.028878611 | 0.977203772 |
| RASSF1 | Memory CD8+ T cell | CellMarker | ENSG00000068028.17 | 0.358490848 | 0.125583187 | 4.799391887 | 1.60808E-06 | 0.971861249 | -0.026693485 | -0.262620855 | 0.79511751 |
| RGS19 | Memory CD8+ T cell | CellMarker | ENSG00000171700.13 | 0.347795764 | -0.218185863 | -8.316674378 | 9.88365E-17 | 0.972747441 | -0.222293496 | -2.54901541 | 0.017726384 |
| RNF166 | Memory CD8+ T cell | CellMarker | ENSG00000158717.10 | 0.434761881 | 0.025762513 | 1.019371068 | 0.308044632 | 0.948629625 | 0.048324565 | 0.291331563 | 0.773338192 |
| S1PR5 | Memory CD8+ T cell | CellMarker | ENSG00000180739.13 | 0.33730812 | 0.137951634 | 5.22937404 | 1.7256E-07 | 0.781297048 | 0.254014106 | 0.686735878 | 0.498929426 |
| SH3BP5 | Memory CD8+ T cell | CellMarker | ENSG00000131370.15 | 0.053113297 | -0.015864243 | -0.566739932 | 0.570900065 | 0.86371251 | -0.33505948 | -1.167420672 | 0.254678775 |
| SLC2A1 | Memory CD8+ T cell | CellMarker | ENSG00000117394.21 | 0.269158211 | 0.137097572 | 5.080923128 | 3.80491E-07 | 0.953084575 | -0.028379789 | -0.182044291 | 0.857099667 |
| SLC9A3R1 | Memory CD8+ T cell | CellMarker | ENSG00000109062.11 | 0.440980513 | -0.06585242 | -2.614876763 | 0.00893563 | 0.954704387 | -0.077430656 | -0.511801121 | 0.61353726 |
| SLCO4C1 | Memory CD8+ T cell | CellMarker | ENSG00000173930.8 | 0.133265873 | -0.079994033 | -2.879987877 | 0.003982998 | 0.697642274 | -0.063794142 | -0.147439615 | 0.884035067 |
| SRPK2 | Memory CD8+ T cell | CellMarker | ENSG00000135250.16 | 0.403005515 | 0.004010745 | 0.156182033 | 0.875891813 | 0.978038077 | -0.002378141 | -0.031488294 | 0.975144418 |
| STK10 | Memory CD8+ T cell | CellMarker | ENSG00000072786.12 | 0.480074362 | 0.008600028 | 0.349200751 | 0.726943908 | 0.97579729 | -0.014494175 | -0.168625595 | 0.867525318 |
| SYNE1 | Memory CD8+ T cell | CellMarker | ENSG00000131018.22 | 0.504563144 | 0.335887218 | 13.95131875 | 6.12107E-44 | 0.952021013 | 0.128842781 | 0.82451536 | 0.417887167 |
| TAGLN2 | Memory CD8+ T cell | CellMarker | ENSG00000158710.14 | 0.810742787 | 0.045697854 | 2.758540155 | 0.005813631 | 0.978448687 | -0.009671847 | -0.131743934 | 0.896301457 |
| TGFBR3 | Memory CD8+ T cell | CellMarker | ENSG00000069702.10 | 0.448177698 | 0.509298354 | 20.60670616 | 5.88028E-93 | 0.86031117 | 0.421098746 | 1.471530382 | 0.15433294 |
| TM4SF19 | Memory CD8+ T cell | CellMarker | ENSG00000145107.15 | 0.031954311 | 0.038151729 | 1.36180562 | 0.173281375 | 0.590801905 | -0.017028735 | -0.034716375 | 0.97259732 |
| TPST2 | Memory CD8+ T cell | CellMarker | ENSG00000128294.15 | 0.4298983 | 0.095107587 | 3.755388516 | 0.000173781 | 0.923510238 | -0.03860337 | -0.180511302 | 0.858289418 |
| TTC16 | Memory CD8+ T cell | CellMarker | ENSG00000167094.15 | 0.086953168 | 0.040856856 | 1.463126406 | 0.143455474 | 0.777700829 | 0.453207225 | 1.242593014 | 0.226200337 |
| TTC38 | Memory CD8+ T cell | CellMarker | ENSG00000075234.16 | 0.260332284 | 0.006723954 | 0.248344848 | 0.803871282 | 0.853621868 | 0.18572643 | 0.611115645 | 0.546953699 |
| USP28 | Memory CD8+ T cell | CellMarker | ENSG00000048028.11 | 0.255286401 | 0.123044703 | 4.541682888 | 5.62776E-06 | 0.960495066 | -0.072507214 | -0.531195933 | 0.600237969 |
| VCL | Memory CD8+ T cell | CellMarker | ENSG00000035403.17 | 0.332565082 | -0.049821164 | -1.883632132 | 0.059635673 | 0.957839216 | 0.104113392 | 0.729553911 | 0.472828855 |
| VEZF1 | Memory CD8+ T cell | CellMarker | ENSG00000136451.8 | 0.389664776 | 0.044954906 | 1.739958697 | 0.081888471 | 0.9725216 | 0.001201168 | 0.012093699 | 0.990452323 |
| WDR37 | Memory CD8+ T cell | CellMarker | ENSG00000047056.15 | 0.231991326 | 0.033672436 | 1.234630704 | 0.216988911 | 0.968811166 | -0.092437948 | -0.833865045 | 0.412702146 |
| ZEB2 | Memory CD8+ T cell | CellMarker | ENSG00000169554.19 | 0.504188347 | 0.027668844 | 1.141026852 | 0.253878467 | 0.955640114 | 0.201655558 | 1.40054231 | 0.174337871 |
| ZNF600 | Memory CD8+ T cell | CellMarker | ENSG00000189190.10 | 0.209015988 | 0.043248012 | 1.577450547 | 0.114714718 | 0.957689677 | -0.001647437 | -0.0113861 | 0.991010929 |
| PTGDS | γδ T cell | Panlaodb | ENSG00000107317.12 | 0.090986065 | -0.067765296 | -2.466074438 | 0.013672312 | 0.762107023 | 0.506605813 | 1.352629386 | 0.188980296 |
| S100B | γδ T cell | Panlaodb | ENSG00000160307.9 | 0.046974686 | -0.052616253 | -1.908836832 | 0.05630378 | 0.44606235 | -0.509785542 | -0.950060497 | 0.351700939 |
| IFITM1 | γδ T cell | Panlaodb | ENSG00000185885.15 | 0.486349896 | 0.156256604 | 6.479312501 | 9.52547E-11 | 0.968952981 | 0.120497978 | 1.102533527 | 0.281335859 |
| CLIC3 | γδ T cell | Panlaodb | ENSG00000169583.12 | 0.095548324 | 0.070183487 | 2.555202266 | 0.010623196 | 0.887094215 | 0.322141758 | 1.244828353 | 0.225403138 |
| STMN1 | γδ T cell | Panlaodb | ENSG00000117632.22 | 0.227226621 | 0.034406416 | 1.279889359 | 0.200605476 | 0.960596141 | 0.129468765 | 0.96258337 | 0.345507146 |
| HMGB2 | γδ T cell | Panlaodb | ENSG00000164104.11 | 0.556440045 | 0.044127914 | 1.920211178 | 0.054851732 | 0.977525276 | 0.08065892 | 1.056731147 | 0.30132171 |
| TYMS | γδ T cell | Panlaodb | ENSG00000176890.15 | 0.017080297 | 0.025476273 | 0.923270593 | 0.355882284 | 0.722739126 | 0.753706092 | 1.94790101 | 0.063405251 |
| MKI67 | γδ T cell | Panlaodb | ENSG00000148773.13 | 0.050899823 | 0.056036197 | 2.033330583 | 0.042038152 | 0.544598268 | 0.413509622 | 0.820637799 | 0.420056946 |
| UBE2C | γδ T cell | Panlaodb | ENSG00000175063.16 | 0.027688381 | -0.013368226 | -0.484573117 | 0.627986846 | 0.570852833 | 0.752147854 | 1.582648319 | 0.126799124 |
| TUBA1B | γδ T cell | Panlaodb | ENSG00000123416.15 | 0.064531641 | -0.038436596 | -1.395691129 | 0.162829888 | 0.971984536 | 0.15853852 | 1.651256434 | 0.111919346 |
| ASPM | γδ T cell | Panlaodb | ENSG00000066279.17 | 0.032003046 | 0.008144829 | 0.29527086 | 0.767791398 | 0.451096309 | 0.540395294 | 1.012587606 | 0.321516847 |
| CENPA | γδ T cell | Panlaodb | ENSG00000115163.14 | 0.018690674 | 0.014025372 | 0.508288832 | 0.611258892 | 0.31807603 | -0.363515462 | -0.631671881 | 0.533666899 |
| TOP2A | γδ T cell | Panlaodb | ENSG00000131747.14 | 0.051426732 | 0.005109566 | 0.185383356 | 0.852931075 | 0.652975663 | 0.674838468 | 1.541583164 | 0.136465189 |
| PCNA | γδ T cell | Panlaodb | ENSG00000132646.10 | 0.182126348 | 0.026223845 | 0.966221176 | 0.33395038 | 0.945518075 | 0.187592682 | 1.112493724 | 0.277119317 |
| AURKB | γδ T cell | Panlaodb | ENSG00000178999.12 | 0.042512035 | -0.016087038 | -0.583427949 | 0.559614742 | 0.596432973 | 0.174548292 | 0.358572731 | 0.723099437 |
| BIRC5 | γδ T cell | Panlaodb | ENSG00000089685.14 | 0.027294902 | -0.030572966 | -1.10824052 | 0.26777719 | 0.511745067 | 0.658979528 | 1.301938389 | 0.205490178 |
| NUSAP1 | γδ T cell | Panlaodb | ENSG00000137804.12 | 0.035538452 | -0.021013378 | -0.761892347 | 0.446137224 | 0.737789613 | 0.483185152 | 1.225815743 | 0.232348427 |
| TROAP | γδ T cell | Panlaodb | ENSG00000135451.12 | 0.02259511 | -0.025308738 | -0.917298034 | 0.359002451 | 0.487571718 | -0.55722482 | -1.070836856 | 0.295062018 |
| TUBB | γδ T cell | Panlaodb | ENSG00000196230.12 | 0.027970154 | 0.069557956 | 2.521922217 | 0.011682664 | 0.907035368 | 0.425811037 | 1.907337255 | 0.068707674 |
| H2AFX | γδ T cell | Panlaodb | ENSG00000188486.3 | 0.139371361 | 0.152939267 | 5.601993076 | 2.15935E-08 | 0.695237746 | 0.858493346 | 2.162115259 | 0.040965255 |
| CENPF | γδ T cell | Panlaodb | ENSG00000117724.12 | 0.062004755 | 0.02243077 | 0.814328169 | 0.415470969 | 0.74385572 | 0.129096322 | 0.321664625 | 0.750535745 |
| CCNB1 | γδ T cell | Panlaodb | ENSG00000134057.14 | 0.055239749 | 0.06218552 | 2.257059045 | 0.024019891 | 0.711394872 | 0.688681058 | 1.721790516 | 0.098178532 |
| H2AFZ | γδ T cell | Panlaodb | ENSG00000164032.11 | 0.448867627 | -0.096311635 | -3.902389236 | 9.56983E-05 | 0.977772203 | 0.041699225 | 0.545612473 | 0.590447781 |
| TRGV2 | γδ T cell | Panlaodb | ENSG00000233306.2 | 0.060616579 | 0.07272184 | 2.64047615 | 0.008288243 | 0.699041126 | 0.867204519 | 2.202919878 | 0.037607918 |
| TRGV10 | γδ T cell | Panlaodb | ENSG00000211694.2 | 0.088649385 | 0.176600212 | 6.433561698 | 1.28743E-10 | 0.832885134 | 0.769620621 | 2.68151295 | 0.013149315 |
| TRGV11 | γδ T cell | Panlaodb | ENSG00000211693.2 | 0.008540149 | 0.079649958 | 2.887013606 | 0.003895186 | 0.199826313 | -0.22837293 | -0.381540825 | 0.706212297 |
| TRGV1 | γδ T cell | Panlaodb | ENSG00000211701.2 | 0.008540149 | -0.009565804 | -0.346621952 | 0.728880615 | 0.40182111 | -0.459224126 | -0.832189366 | 0.413636023 |
| TRGV3 | γδ T cell | Panlaodb | ENSG00000211699.2 | 0.05029369 | 0.247595897 | 9.008922621 | 2.34921E-19 | 0.831628578 | 0.597908058 | 1.95930712 | 0.061980559 |
| TRGV4 | γδ T cell | Panlaodb | ENSG00000211698.2 | 0.078373199 | 0.347341942 | 12.69726333 | 9.80788E-37 | 0.769483644 | 1.128023799 | 3.700856529 | 0.001140025 |
| TRGV5 | γδ T cell | Panlaodb | ENSG00000211697.4 | 0.050642655 | 0.14183308 | 5.150646907 | 2.6315E-07 | 0.686749707 | 0.935797693 | 2.366822941 | 0.026492584 |
| TRGV8 | γδ T cell | Panlaodb | ENSG00000211696.2 | 0.05081478 | 0.17158683 | 6.233970822 | 4.67943E-10 | 0.676587969 | 1.172491501 | 3.12732067 | 0.004633146 |
| TRGV9 | γδ T cell | Panlaodb | ENSG00000211695.2 | 0.096673458 | 0.310120013 | 11.34151517 | 1.10446E-29 | 0.815814934 | 0.882012236 | 3.01802271 | 0.006011111 |
| TRGJ1 | γδ T cell | Panlaodb | ENSG00000211690.1 | 0.014791971 | 0.08644903 | 3.133848744 | 0.001728897 | 0.638096042 | 1.778843785 | 6.123188911 | 2.69508E-06 |
| TRGJ2 | γδ T cell | Panlaodb | ENSG00000211687.1 | 0.019096353 | 0.098305718 | 3.564291248 | 0.000366068 | 0.723388986 | 1.012310501 | 2.807425047 | 0.009846594 |
| TRGJP1 | γδ T cell | Panlaodb | ENSG00000211692.1 | 0.023586171 | -0.026418823 | -0.95755663 | 0.338303083 | 0.657242072 | 0.948871849 | 2.297085187 | 0.030791653 |
| TRGJP2 | γδ T cell | Panlaodb | ENSG00000211688.1 | 0.035211935 | 0.176939187 | 6.424710204 | 1.36437E-10 | 0.715424462 | 1.151674809 | 3.301445182 | 0.003044321 |
| TRGJP | γδ T cell | Panlaodb | ENSG00000211691.2 | 0.029583943 | 0.0441262 | 1.599712146 | 0.109685253 | 0.565229151 | 1.376697269 | 3.319035135 | 0.002916941 |
| TRGC1 | γδ T cell | Panlaodb | ENSG00000211689.7 | 0.316333073 | 1.791466582 | 84.01867174 | 0 | 0.855469019 | 0.754879258 | 2.882799995 | 0.00826324 |
| TRGC2 | γδ T cell | Panlaodb | ENSG00000227191.8 | 0.357756151 | 1.169586662 | 49.14101152 | 0 | 0.871963279 | 0.553097326 | 2.109990147 | 0.045645998 |
| TRDD1 | γδ T cell | Panlaodb | ENSG00000223997.1 | NA | NA | NA | NA | 0.199826313 | 1.598610508 | 3.18264562 | 0.004057033 |
| TRDD2 | γδ T cell | Panlaodb | ENSG00000237235.2 | NA | NA | NA | NA | 0.342626766 | 2.741014129 | 24.56264238 | 2.51147E-18 |
| TRDD3 | γδ T cell | Panlaodb | ENSG00000228985.1 | 0.008540149 | 0.079649958 | 2.887013606 | 0.003895186 | 0.395145343 | 0.231880444 | 0.414321591 | 0.682376658 |
| TRDJ1 | γδ T cell | Panlaodb | ENSG00000211825.1 | 0.034160595 | -0.015959275 | -0.578610589 | 0.562861362 | 0.58042452 | 1.156209545 | 2.659867173 | 0.013813045 |
| TRDJ2 | γδ T cell | Panlaodb | ENSG00000211827.1 | 0.019096353 | 0.098305718 | 3.564291248 | 0.000366068 | 0.595168136 | 1.59152935 | 4.387023194 | 0.000203597 |
| TRDJ3 | γδ T cell | Panlaodb | ENSG00000211828.1 | NA | NA | NA | NA | 0.403337388 | 2.554129309 | 13.31151609 | 1.8077E-12 |
| TRDJ4 | γδ T cell | Panlaodb | ENSG00000211826.1 | 0.014791971 | 0.034940286 | 1.266238881 | 0.205448856 | 0.442583921 | 0.77538586 | 1.479497952 | 0.152220251 |
| TRDV1 | γδ T cell | Panlaodb | ENSG00000211804.3 | 0.029583943 | 0.224406805 | 8.154186668 | 3.81533E-16 | 0.605329196 | 1.691075515 | 5.029390725 | 4.0292E-05 |
| TRDV2 | γδ T cell | Panlaodb | ENSG00000211821.2 | 0.060616579 | 0.565342382 | 20.84114048 | 5.22533E-95 | 0.401893763 | 2.602731837 | 15.94171098 | 3.85547E-14 |
| TRDV3 | γδ T cell | Panlaodb | ENSG00000256590.2 | 0.052919613 | 0.466031746 | 17.0868781 | 8.55728E-65 | 0.448490999 | 2.449310901 | 11.65943084 | 2.77758E-11 |
| CD24 | Naive B cell | CellMarker | ENSG00000272398.5 | 0.136084724 | 1.130778698 | 28.6033621 | 6.9879E-175 | 0.671224572 | 1.266560009 | 3.466604691 | 0.002033548 |
| IL2RA | Naive B cell | Panlaodb | ENSG00000134460.17 | 0.098197584 | 0.111155533 | 2.720801024 | 0.006520521 | 0.67739192 | 1.034148963 | 2.644703256 | 0.01429578 |
| ABCB4 | Naive B cell | Panlaodb | ENSG00000005471.17 | 0.057034632 | 0.766323494 | 18.93324945 | 6.00347E-79 | 0.444814644 | 2.192654441 | 7.088678598 | 2.74117E-07 |
| BCL7A | Naive B cell | Panlaodb | ENSG00000110987.8 | 0.072868586 | 0.633768347 | 15.61088183 | 1.78532E-54 | 0.658338163 | 1.400158243 | 3.959627629 | 0.000597249 |
| BEND5 | Naive B cell | Panlaodb | ENSG00000162373.12 | 0.075909149 | 0.492357352 | 12.08836203 | 1.79326E-33 | 0.665672258 | 1.22925153 | 3.281776334 | 0.003192791 |
| BRAF | Naive B cell | Panlaodb | ENSG00000157764.13 | 0.291701446 | 0.126149093 | 3.211555129 | 0.0013232 | 0.978130589 | 0.064388775 | 0.882575293 | 0.386357793 |
| EAF2 | Naive B cell | Panlaodb | ENSG00000145088.8 | 0.185062566 | 1.202491372 | 30.79725118 | 1.627E-201 | 0.942135684 | 0.552224495 | 3.971026861 | 0.000580412 |
| FCER2 | Naive B cell | Panlaodb | ENSG00000104921.14 | 0.119804654 | 1.824761353 | 48.3932309 | 0 | 0.563058526 | 1.761208217 | 5.047245672 | 3.85116E-05 |
| IL4R | Naive B cell | Panlaodb | ENSG00000077238.13 | 0.248292673 | 0.901411191 | 23.08633461 | 9.5738E-116 | 0.954848175 | 0.316089533 | 2.312407262 | 0.029794599 |
| MEP1A | Naive B cell | Panlaodb | ENSG00000112818.9 | NA | NA | NA | NA | 0.199826313 | -0.22837293 | -0.381641702 | 0.706138036 |
| MICAL3 | Naive B cell | Panlaodb | ENSG00000243156.7 | 0.073500236 | 0.170361378 | 4.162690159 | 3.16421E-05 | 0.880734284 | 0.552159038 | 2.205899001 | 0.037371542 |
| P2RY14 | Naive B cell | Panlaodb | ENSG00000174944.8 | 0.099011569 | 0.388541265 | 9.539883036 | 1.6647E-21 | 0.529702132 | 2.207835045 | 8.951206509 | 4.6866E-09 |
| PSG2 | Naive B cell | Panlaodb | ENSG00000242221.8 | NA | NA | NA | NA | 0.199826313 | -0.22837293 | -0.381641702 | 0.706138036 |
| TCL1A | Naive B cell | Panlaodb | ENSG00000100721.10 | 0.115714905 | 1.940376816 | 52.00734875 | 0 | 0.56190596 | 1.978833594 | 6.677721685 | 7.15128E-07 |
| UGT1A8 | Naive B cell | Panlaodb | ENSG00000242366.3 | NA | NA | NA | NA | 0.199826313 | -0.22837293 | -0.381641702 | 0.706138036 |
| ZNF286A | Naive B cell | Panlaodb | ENSG00000187607.15 | 0.080916723 | 0.015478916 | 0.378200385 | 0.705287532 | 0.954828606 | 0.056932497 | 0.377118221 | 0.709451945 |
| MITF | Naive B cell | Panlaodb | ENSG00000187098.15 | 0.086341854 | -0.073290186 | -1.79172628 | 0.073198632 | 0.554123185 | -0.261910018 | -0.519667906 | 0.608130022 |
| CD52 | Naive B cell | Panlaodb | ENSG00000169442.8 | 0.57849017 | 0.396219631 | 11.85893214 | 2.77081E-32 | 0.975844791 | 0.088882862 | 1.070643237 | 0.295145972 |
| HLA-DQB1 | Naive B cell | Panlaodb | ENSG00000179344.16 | 0.335760797 | 1.006275623 | 26.65827372 | 9.8656E-153 | 0.945289734 | 0.375272683 | 2.423305681 | 0.023421634 |
| TNFRSF13C | Naive B cell | Panlaodb | ENSG00000159958.6 | 0.222087275 | 1.981946977 | 54.54534276 | 0 | 0.88616897 | 0.610815865 | 2.579973739 | 0.016544009 |
| LINC00926 | Naive B cell | Panlaodb | ENSG00000247982.6 | 0.16568521 | 1.943513447 | 52.55294827 | 0 | 0.702299655 | 1.195266271 | 3.388870624 | 0.002460077 |
| STAG3 | Naive B cell | Panlaodb | ENSG00000066923.17 | 0.065379386 | 0.171786625 | 4.195203209 | 2.74324E-05 | 0.820029561 | 0.614419251 | 1.947501366 | 0.063454175 |
| BIRC3 | Naive B cell | Panlaodb | ENSG00000023445.13 | 0.447076118 | 0.976247512 | 27.2471432 | 2.726E-159 | 0.975578709 | 0.14177942 | 1.748295973 | 0.093397395 |
| CD74 | Naive B cell | Panlaodb | ENSG00000019582.14 | 0.669612355 | 1.002124717 | 34.06551821 | 2.4664E-244 | 0.970316499 | 0.224726193 | 2.33629303 | 0.028300792 |
| CD84 | Memory B cell | Panlaodb | ENSG00000066294.14 | 0.422185294 | -0.052088159 | -1.040864746 | 0.297956483 | 0.939647959 | -0.669341648 | -5.520072507 | 1.19464E-05 |
| PAX5 | Memory B cell | Panlaodb | ENSG00000196092.12 | 0.188151011 | 2.364408077 | 46.99766971 | 0 | 0.616207838 | 1.361658052 | 3.511900481 | 0.001825083 |
| AIM2 | Memory B cell | Panlaodb | ENSG00000163568.14 | 0.097737689 | 1.385055281 | 25.83705744 | 8.4089E-144 | 0.657556837 | 1.388010986 | 3.897722419 | 0.000700464 |
| ALOX5 | Memory B cell | Panlaodb | ENSG00000012779.10 | 0.115877435 | 0.238237996 | 4.352217146 | 1.35733E-05 | 0.529167843 | 1.571051959 | 3.891829537 | 0.000710862 |
| CCR6 | Memory B cell | Panlaodb | ENSG00000112486.16 | 0.038851578 | 0.035240591 | 0.639546687 | 0.522477923 | 0.723210421 | -0.261755916 | -0.633906921 | 0.532249529 |
| CLCA3P | Memory B cell | Panlaodb | ENSG00000153923.10 | NA | NA | NA | NA | 0.275627857 | -0.315003265 | -0.538073834 | 0.595574159 |
| DENND5B | Memory B cell | Panlaodb | ENSG00000170456.15 | 0.114939984 | 1.27949419 | 23.82943719 | 4.8591E-123 | 0.654238225 | 1.147532951 | 2.920508301 | 0.007578863 |
| GNG7 | Memory B cell | Panlaodb | ENSG00000176533.12 | 0.200677026 | 1.85951169 | 35.98529713 | 3.4975E-271 | 0.83601188 | 0.754012026 | 2.649115526 | 0.014173646 |
| IFNA10 | Memory B cell | Panlaodb | ENSG00000186803.3 | NA | NA | NA | NA | NA | NA | NA | NA |
| IL7 | Memory B cell | Panlaodb | ENSG00000104432.13 | 0.060456025 | 0.607166795 | 11.07918203 | 2.08337E-28 | 0.646866665 | 0.05735632 | 0.124023003 | 0.902349317 |
| MBL2 | Memory B cell | Panlaodb | ENSG00000165471.6 | NA | NA | NA | NA | 0.199826313 | -0.22837293 | -0.381304056 | 0.706395322 |
| NPIPB15 | Memory B cell | Panlaodb | ENSG00000196436.8 | 0.15358474 | 0.028089117 | 0.515429737 | 0.606260978 | 0.953070713 | 0.136345896 | 0.897624285 | 0.378469248 |
| SIT1 | Memory B cell | Panlaodb | ENSG00000137078.8 | 0.183891918 | 0.293731759 | 5.423764156 | 5.93357E-08 | 0.802080647 | -0.548848647 | -1.626018064 | 0.117251368 |
| SP140 | Memory B cell | Panlaodb | ENSG00000079263.18 | 0.277097374 | 0.972310092 | 18.56951853 | 4.73516E-76 | 0.968404587 | 0.193661289 | 1.860618114 | 0.075327534 |
| TMEM156 | Memory B cell | Panlaodb | ENSG00000121895.7 | 0.17453425 | 0.850055631 | 15.79303427 | 1.06257E-55 | 0.946641609 | 0.186929366 | 1.135120839 | 0.267743451 |
| TNFRSF13B | Memory B cell | Panlaodb | ENSG00000240505.8 | 0.09107957 | 2.164107177 | 41.81882249 | 0 | 0.473752403 | 1.532819933 | 3.520547641 | 0.001786669 |
| TNFRSF17 | Memory B cell | Panlaodb | ENSG00000048462.10 | 0.044538882 | 0.174993153 | 3.17763466 | 0.001488077 | 0.468930134 | -0.535920153 | -1.014821399 | 0.320501954 |
| TRAF4 | Memory B cell | Panlaodb | ENSG00000076604.14 | 0.071469031 | 0.573213342 | 10.46220072 | 1.60354E-25 | 0.714757423 | -0.216206478 | -0.5151439 | 0.611253213 |
| ZBTB32 | Memory B cell | Panlaodb | ENSG00000011590.13 | 0.059887069 | 1.022695374 | 18.81451882 | 5.36846E-78 | 0.467629762 | 1.417021215 | 3.119761705 | 0.004728133 |
| NCAM1 | NK cell | CellMarker | ENSG00000149294.16 | 0.153814843 | 0.900283215 | 36.58912325 | 7.0833E-280 | 0.575006967 | 1.73683208 | 5.008905182 | 4.24284E-05 |
| KLRB1 | NK cell | CellMarker/Panlaodb | ENSG00000111796.3 | 0.411313692 | 0.732950952 | 31.92145205 | 7.9916E-216 | 0.84272151 | 0.507581462 | 1.690746491 | 0.104040166 |
| KLRC1 | NK cell | CellMarker/Panlaodb | ENSG00000134545.13 | 0.164625844 | 0.37654869 | 14.7539839 | 6.81216E-49 | 0.74578666 | 0.910944892 | 2.571366259 | 0.016868165 |
| KLRK1 | NK cell | CellMarker/Panlaodb | ENSG00000213809.8 | 0.035211935 | 0.017827058 | 0.684150521 | 0.493891565 | 0.848263785 | 0.394547299 | 1.309233359 | 0.203048488 |
| DPP4 | NK cell | CellMarker/Panlaodb | ENSG00000197635.9 | 0.17966812 | -0.194330315 | -7.590874718 | 3.3801E-14 | 0.763581336 | -0.818494864 | -2.344530969 | 0.027803522 |
| ITGA1 | NK cell | CellMarker | ENSG00000213949.8 | 0.069545464 | -0.070694454 | -2.718574579 | 0.006564547 | 0.652854234 | -0.0568121 | -0.123736911 | 0.902570468 |
| CXCR6 | NK cell | CellMarker | ENSG00000172215.5 | 0.086653229 | -0.084643236 | -3.259693465 | 0.001118017 | 0.552821777 | -0.392164484 | -0.782587917 | 0.441644468 |
| CD160 | NK cell | CellMarker | ENSG00000117281.15 | 0.181259038 | 0.741617034 | 29.83147945 | 1.4111E-189 | 0.733290671 | 1.044545417 | 2.995471439 | 0.006341013 |
| ID2 | NK cell | CellMarker | ENSG00000115738.9 | 0.483425549 | 0.212240188 | 9.313508704 | 1.42022E-20 | 0.936087127 | 0.211856233 | 1.134826878 | 0.267832685 |
| IL2RB | NK cell | Panlaodb | ENSG00000100385.13 | 0.451056642 | 1.488338568 | 76.04622169 | 0 | 0.86459785 | 0.630853679 | 2.389026271 | 0.025244755 |
| CD244 | NK cell | Panlaodb | ENSG00000122223.12 | 0.203947086 | 0.435260947 | 17.22984087 | 7.68702E-66 | 0.821931268 | 0.785381404 | 2.642771529 | 0.014359835 |
| SLAMF7 | NK cell | Panlaodb | ENSG00000026751.16 | 0.371448881 | 0.588608132 | 24.82941046 | 3.4461E-133 | 0.836734175 | 0.489597643 | 1.590322658 | 0.125058011 |
| SIGLEC7 | NK cell | Panlaodb | ENSG00000168995.13 | 0.06652089 | 0.085366099 | 3.282505582 | 0.001031436 | 0.400303693 | 2.678618556 | 26.59814057 | 4.08742E-19 |
| NCR1 | NK cell | Panlaodb | ENSG00000189430.12 | 0.052227357 | 0.253429476 | 9.766342463 | 1.85523E-22 | 0.54699658 | 1.390471988 | 3.293381732 | 0.003104697 |
| SLAMF6 | NK cell | Panlaodb | ENSG00000162739.13 | 0.378095648 | 0.208053437 | 8.63517614 | 6.49743E-18 | 0.959715174 | 0.080430426 | 0.581365204 | 0.566501188 |
| KIT | NK cell | Panlaodb | ENSG00000157404.15 | 0.047323771 | -0.007615186 | -0.292389504 | 0.769993211 | 0.503902468 | -0.354700777 | -0.679724576 | 0.503286276 |
| IL32 | NK cell | Panlaodb | ENSG00000008517.16 | 0.501974872 | -0.109841797 | -4.866779336 | 1.14681E-06 | 0.91004362 | 0.157925114 | 0.677687452 | 0.504554232 |
| GZMM | NK cell | Panlaodb | ENSG00000197540.7 | 0.355119946 | 0.519907281 | 21.67298198 | 1.8414E-102 | 0.684425678 | 0.249168044 | 0.568685719 | 0.574936933 |
| HMBOX1 | NK cell | Panlaodb | ENSG00000147421.17 | 0.272245754 | 0.009839195 | 0.392018447 | 0.695050613 | 0.955829133 | 0.046539015 | 0.312325457 | 0.757533654 |
| AHR | NK cell | Panlaodb | ENSG00000106546.13 | 0.320809811 | -0.304685079 | -12.39874576 | 4.07563E-35 | 0.948017559 | -0.434182289 | -3.064712791 | 0.005380399 |
| CCL4 | NK cell | Panlaodb | ENSG00000275302.1 | 0.295791354 | 0.932603275 | 39.4737635 | 5e-323 | 0.868049609 | 0.611901147 | 2.339464606 | 0.028109494 |
| SEMA6D | NK cell | Panlaodb | ENSG00000137872.16 | NA | NA | NA | NA | 0.346109328 | -0.395553517 | -0.696074036 | 0.49317515 |
| FHL2 | NK cell | Panlaodb | ENSG00000115641.18 | 0.055603088 | -0.01639063 | -0.629601028 | 0.528966032 | 0.634166657 | -0.257668943 | -0.552574879 | 0.585746022 |
| LAT | NK cell | Panlaodb | ENSG00000213658.11 | 0.087271489 | 0.003987164 | 0.153499094 | 0.87800694 | 0.826054327 | 0.174230423 | 0.524100787 | 0.605091915 |
| PCDH15 | NK cell | Panlaodb | ENSG00000150275.18 | NA | NA | NA | NA | 0.431512624 | -0.493157285 | -0.91022152 | 0.371900682 |
| CMA1 | NK cell | Panlaodb | ENSG00000092009.10 | NA | NA | NA | NA | 0.199826313 | -0.22837293 | -0.38155003 | 0.706205797 |
| DUSP2 | NK cell | Panlaodb | ENSG00000158050.4 | 0.55287361 | 0.357414087 | 16.57626967 | 4.00802E-61 | 0.953853128 | 0.222774734 | 1.513875121 | 0.143324131 |
| DOK2 | NK cell | Panlaodb | ENSG00000147443.12 | 0.272081741 | 0.185646189 | 7.410864763 | 1.32738E-13 | 0.894822379 | 0.467747294 | 1.963792874 | 0.061428871 |
| HSD11B1 | NK cell | Panlaodb | ENSG00000117594.9 | 0.014791971 | -0.01685068 | -0.646351828 | 0.518062236 | 0.343441799 | -0.392504913 | -0.68984073 | 0.497016351 |
| KLRC2 | NK cell | Panlaodb | ENSG00000205809.9 | 0.149476825 | 0.350544971 | 13.68720484 | 2.28317E-42 | 0.711600715 | 0.918164348 | 2.417657316 | 0.023714838 |
| CHSY1 | NK cell | Panlaodb | ENSG00000131873.6 | 0.287817734 | 0.004850477 | 0.194161047 | 0.846052629 | 0.88149028 | 0.074552133 | 0.272361054 | 0.787711073 |
| XCL2 | NK cell | Panlaodb | ENSG00000143185.3 | 0.143973611 | 0.45202273 | 17.71427053 | 1.89431E-69 | 0.760185452 | 0.878245891 | 2.539647335 | 0.018108868 |
| IL18R1 | NK cell | Panlaodb | ENSG00000115604.10 | 0.177657909 | 0.340330965 | 13.34645702 | 2.20535E-40 | 0.824449125 | 0.559386097 | 1.772679319 | 0.089179912 |

| | | | | | | | | | | | |
|---|---|---|---|---|---|---|---|---|---|---|---|
| SERPINB9 | NK cell | Panlaodb | ENSG00000170542.5 | 0.383974716 | -0.072785734 | -3.021654662 | 0.002518561 | 0.963126929 | 0.057993515 | 0.448333642 | 0.657997878 |
| DOCK2 | NK cell | Panlaodb | ENSG00000134516.16 | 0.413218423 | -0.075143879 | -3.162671476 | 0.00156666 | 0.976302297 | 0.03431247 | 0.412043884 | 0.684022581 |
| SH2D2A | NK cell | Panlaodb | ENSG00000027869.11 | 0.324403244 | 0.69183712 | 28.86492327 | 5.7716E-178 | 0.873759827 | 0.523710038 | 1.996601411 | 0.05751804 |
| XCL1 | NK cell | Panlaodb | ENSG00000143184.4 | 0.102962458 | 0.40301748 | 15.67581404 | 6.5552E-55 | 0.761405458 | 0.836050188 | 2.394280145 | 0.024957371 |
| GZMK | NK cell | Panlaodb | ENSG00000113088.5 | 0.182809139 | -0.151140089 | -5.902391676 | 3.66643E-09 | 0.815314413 | -0.032726178 | -0.09503746 | 0.9250867 |
| IFNG | NK cell | Panlaodb | ENSG00000111537.4 | 0.123984577 | 0.429476492 | 16.76596003 | 1.78541E-62 | 0.820017142 | 0.781494165 | 2.608012581 | 0.015533691 |
| CSF2 | NK cell | Panlaodb | ENSG00000164400.5 | 0.012077594 | 0.042548744 | 1.632141539 | 0.102672398 | 0.199826313 | -0.22837293 | -0.38155003 | 0.706205797 |
| IL2RG | NK cell | Panlaodb | ENSG00000147168.12 | 0.653169577 | 0.30058571 | 15.29531248 | 2.20005E-52 | 0.9775594 | 0.075826124 | 0.996481324 | 0.329115348 |
| TGFB1 | NK cell | Panlaodb | ENSG00000105329.9 | 0.271266218 | 0.237877048 | 9.505722499 | 2.30795E-21 | 0.923087189 | 0.314219305 | 1.534572274 | 0.138175409 |
| KIR2DL1 | NK cell | Panlaodb | ENSG00000125498.19 | 0.074351735 | 0.419964363 | 16.3053154 | 3.21758E-59 | 0.592226061 | 1.517644739 | 4.018166017 | 0.000515791 |
| KIR3DL1 | NK cell | Panlaodb | ENSG00000167633.18 | 0.050366919 | 0.284547789 | 10.97434314 | 6.61481E-28 | 0.668702871 | 1.255430205 | 3.402854654 | 0.002377858 |
| KLRG1 | NK cell | Panlaodb | ENSG00000139187.9 | 0.34156565 | -0.079556949 | -3.245557505 | 0.001174992 | 0.853249591 | 0.336970722 | 1.127529561 | 0.270841974 |
| NCR3 | NK cell | Panlaodb | ENSG00000204475.9 | 0.188455369 | 0.429079779 | 16.92589784 | 1.26209E-63 | 0.940911709 | 0.282118989 | 1.632158015 | 0.11590761 |
| ADAMTS14 | NK cell | Panlaodb | ENSG00000138316.10 | 0.014791971 | 0.02912401 | 1.117160958 | 0.263944882 | 0.504797052 | 1.564865856 | 3.755633616 | 0.000994859 |
| ITGA2 | NK cell | Panlaodb | ENSG00000164171.10 | 0.032527096 | -0.037054133 | -1.421981211 | 0.155054215 | 0.574652841 | -0.394360532 | -0.802458803 | 0.430287704 |
| STYK1 | NK cell | Panlaodb | ENSG00000060140.8 | 0.053285293 | -0.013762596 | -0.528583842 | 0.597102638 | 0.503272524 | -0.575168599 | -1.119688691 | 0.274102823 |
| CLEC2D | NK cell | Panlaodb | ENSG00000069493.14 | 0.482218127 | -0.133076535 | -5.824211755 | 5.86554E-09 | 0.976622623 | -0.124737467 | -1.599258413 | 0.123054706 |
| ZBTB16 | NK cell | Panlaodb | ENSG00000109906.13 | 0.163741992 | 0.421985066 | 16.5651215 | 4.8071E-61 | 0.806829798 | 0.82306663 | 2.666583603 | 0.013604265 |
| LAIR2 | NK cell | Panlaodb | ENSG00000167618.9 | 0.188061944 | 0.634835211 | 25.35393437 | 1.1516E-138 | 0.805303965 | 0.72053026 | 2.245925651 | 0.034341585 |
| CD40 | Monocytes | Panlaodb/CellMarker | ENSG00000101017.13 | 0.146157265 | 0.131731677 | 4.669549869 | 3.04703E-06 | | | | |
| S100A12 | Monocytes | Panlaodb/CellMarker | ENSG00000163221.8 | 0.317279355 | 1.910274813 | 88.22945292 | 0 | | | | |
| RGS1 | Monocytes | Panlaodb | ENSG00000090104.11 | 0.440319692 | -0.383454657 | -15.07572067 | 5.94712E-51 | | | | |
| APOBEC3A | Monocytes | Panlaodb | ENSG00000128383.12 | 0.347338726 | 1.456581637 | 61.37597513 | 0 | | | | |
| TET2 | Monocytes | Panlaodb | ENSG00000168769.13 | 0.371049901 | 0.981814937 | 39.01578888 | 5.79422324e-316 | | | | |
| DYSF | Monocytes | Panlaodb | ENSG00000135636.13 | 0.211755478 | 0.668600682 | 24.4832458 | 1.2662E-129 | | | | |
| CMKLR1 | Monocytes | Panlaodb | ENSG00000174600.13 | 0.153330596 | 0.023364747 | 0.828491217 | 0.407406683 | | | | |
| MEFV | Monocytes | Panlaodb | ENSG00000103313.12 | 0.242950919 | 0.873056575 | 32.73152694 | 2.2032E-226 | | | | |
| HCK | Monocytes | Panlaodb | ENSG00000101336.13 | 0.386769208 | 1.606740615 | 71.39287724 | 0 | | | | |
| PADI4 | Monocytes | Panlaodb | ENSG00000159339.13 | 0.158142075 | 0.64612642 | 23.37710399 | 1.4444E-118 | | | | |
| GHSR | Monocytes | Panlaodb | ENSG00000121853.3 | 0.012077594 | -0.013410854 | -0.469978543 | 0.638377768 | | | | |
| SELE | Monocytes | Panlaodb | ENSG00000007908.15 | 0.008540149 | -0.009482906 | -0.332311651 | 0.739658986 | | | | |
| TLR4 | Monocytes | Panlaodb | ENSG00000136869.14 | 0.31168476 | 1.6176283 | 69.20472009 | 0 | | | | |
| AR | Monocytes | Panlaodb | ENSG00000169083.16 | 0.020383342 | 0.141524735 | 4.964731771 | 6.9617E-07 | | | | |
| CCR2 | Monocytes | Panlaodb | ENSG00000121807.5 | 0.146191188 | 0.998291057 | 37.07704287 | 5.9869E-287 | | | | |
| TNFRSF14 | Monocytes | Panlaodb | ENSG00000157873.17 | 0.147515881 | 0.108830611 | 3.857581131 | 0.000115035 | | | | |
| ADA2 | Monocytes | Panlaodb | ENSG00000093072.16 | 0.43911735 | 2.08929808 | 112.9083483 | 0 | | | | |
| MGMT | Monocytes | Panlaodb | ENSG00000170430.9 | 0.147559887 | 0.251740363 | 8.944168244 | 4.21808E-19 | | | | |
| ACE | Monocytes | Panlaodb | ENSG00000159640.15 | 0.036232782 | -0.017749949 | -0.622405073 | 0.53368582 | | | | |
| ICAM1 | Monocytes | Panlaodb | ENSG00000090339.8 | 0.179410121 | 0.765159371 | 28.0151858 | 5.0589E-168 | | | | |
| CD163 | Monocytes | Panlaodb | ENSG00000177575.12 | 0.188046652 | 1.677803129 | 69.54868168 | 0 | | | | |
| ACP5 | Monocytes | Panlaodb | ENSG00000102575.11 | 0.119703148 | 0.291407249 | 10.32447998 | 6.72708E-25 | | | | |
| MRC1 | Monocytes | Panlaodb | ENSG00000260314.2 | 0.017237464 | 0.127216073 | 4.461752493 | 8.19347E-06 | | | | |
| TNF | Monocytes | Panlaodb | ENSG00000232810.3 | 0.019096353 | -0.021204422 | -0.743190989 | 0.457378698 | | | | |
| PLAU | Monocytes | Panlaodb | ENSG00000122861.15 | 0.036232782 | 0.094663116 | 3.320656889 | 0.000900386 | | | | |
| GBP1 | Monocytes | Panlaodb | ENSG00000117228.9 | 0.254912017 | 0.366002544 | 13.34554614 | 2.23612E-40 | | | | |
| OAS1 | Monocytes | Panlaodb | ENSG00000089127.12 | 0.170325737 | 1.008186147 | 37.64277106 | 2.7206E-295 | | | | |
| IRF7 | Monocytes | Panlaodb | ENSG00000185507.20 | 0.135961294 | 0.440419589 | 15.71537141 | 3.56505E-55 | | | | |
| PLSCR1 | Monocytes | Panlaodb | ENSG00000188313.12 | 0.279262642 | 1.396031061 | 56.51378182 | 0 | | | | |
| MX1 | Monocytes | Panlaodb | ENSG00000157601.13 | 0.206448761 | 0.411824851 | 14.86398567 | 1.3642E-49 | | | | |
| IL1RN | Monocytes | Panlaodb | ENSG00000136689.18 | 0.240628957 | 0.923999419 | 34.7815129 | 3.6107E-254 | | | | |
| HLA-DRA | Monocytes | Panlaodb | ENSG00000204287.13 | 0.326304128 | 1.783353211 | 79.91238112 | 0 | | | | |
| IFIT1 | Monocytes | Panlaodb | ENSG00000185745.9 | 0.091375492 | 0.032129283 | 1.130617255 | 0.258235854 | | | | |
| IDO1 | Monocytes | Panlaodb | ENSG00000131203.12 | 0.030365461 | 0.06395304 | 2.242465692 | 0.02494709 | | | | |
| IFIT3 | Monocytes | Panlaodb | ENSG00000119917.13 | 0.183962033 | 0.028654481 | 1.021452145 | 0.307058161 | | | | |
| TNFSF10 | Monocytes | Panlaodb | ENSG00000121858.10 | 0.27606138 | 0.773945725 | 29.05825668 | 3.0333E-180 | | | | |
| CXCL10 | Monocytes | Panlaodb | ENSG00000169245.5 | 0.018632868 | 0.159477209 | 5.595667161 | 2.23954E-08 | | | | |
| S100A9 | Monocytes | Panlaodb | ENSG00000163220.10 | 0.516049424 | 1.770430281 | 91.78063927 | 0 | | | | |
| S100A8 | Monocytes | Panlaodb | ENSG00000143546.9 | 0.50623243 | 1.679121544 | 83.64132861 | 0 | | | | |
| CLEC7A | Monocytes | Panlaodb | ENSG00000172243.17 | 0.421971143 | 1.678460742 | 77.71797263 | 0 | | | | |
| MS4A6A | Monocytes | Panlaodb | ENSG00000110077.14 | 0.313807213 | 2.592615773 | 164.599493 | 0 | | | | |
| ZFP36L2 | Monocytes | Panlaodb | ENSG00000152518.7 | 0.761854855 | -0.049866558 | -2.685343089 | 0.007254194 | | | | |
| LTA4H | Monocytes | Panlaodb | ENSG00000111144.9 | 0.390298379 | 1.970346709 | 97.29458281 | 0 | | | | |
| CLEC12A | Monocytes | Panlaodb | ENSG00000172322.13 | 0.350205646 | 1.859601502 | 86.25238763 | 0 | | | | |
| CD48 | Monocytes | Panlaodb | ENSG00000117091.9 | 0.500486404 | 0.534521947 | 21.98473253 | 2.5558E-105 | | | | |
| PRTN3 | Monocytes | Panlaodb | ENSG00000196415.9 | | 2.604975516 | 167.0812105 | 0 | | | | |
| VCAN | Monocytes | Panlaodb | ENSG00000038427.15 | 0.314521817 | 0.841790928 | 34.00967237 | 1.5361E-243 | | | | |
| IFITM3 | Monocytes | Panlaodb | ENSG00000142089.15 | 0.43131902 | 0.057042401 | 2.000784819 | 0.045435119 | | | | |
| FN1 | Monocytes | Panlaodb | ENSG00000115414.18 | 0.040444807 | 0.885621165 | 32.40512242 | 4.3109E-222 | | | | |
| ADGRE1 | Monocytes | Panlaodb | ENSG00000174837.14 | 0.114938765 | 1.869887486 | 81.82264681 | 0 | | | | |
| CSF1R | Monocytes | Panlaodb | ENSG00000182578.13 | 0.219541231 | 0.102241452 | 4.514272763 | 6.40597E-06 | | | | |
| ITGAL | Monocytes | Panlaodb | ENSG00000005844.17 | 0.609767177 | 0.317957062 | 12.89673457 | 7.76303E-38 | | | | |
| SPN | Monocytes | Panlaodb | ENSG00000197471.11 | 0.496368744 | 2.017420439 | 140.1951067 | 0 | | | | |
| PSAP | Monocytes | Panlaodb | ENSG00000197746.13 | 0.62178335 | 2.638452672 | 180.9088698 | 0 | | | | |
| FCN1 | Monocytes | Panlaodb | ENSG00000085265.10 | 0.347522293 | 2.681390762 | 185.9592877 | 0 | | | | |
| LYZ | Monocytes | Panlaodb | ENSG00000090382.6 | 0.325952899 | 0.105223111 | 3.759623524 | 0.000170871 | | | | |
| RHOC | Monocytes | Panlaodb | ENSG00000155366.16 | 0.193468583 | 1.714915091 | 76.22818669 | 0 | | | | |
| PILRA | Monocytes | Panlaodb | ENSG00000085514.15 | 0.34335416 | 0.770401786 | 31.73537829 | 2.1116E-213 | | | | |
| NFKBIZ | Monocytes | Panlaodb | ENSG00000144802.11 | 0.474534038 | 1.396953707 | 58.15191372 | 0 | | | | |
| NAAA | Monocytes | Panlaodb | ENSG00000138744.14 | 0.34457247 | 0.1257459 | 4.443609505 | 8.91505E-06 | | | | |
| LY6E | Monocytes | Panlaodb | ENSG00000160932.10 | 0.124078181 | 1.571662374 | 76.51945795 | 0 | | | | |
| LYN | Monocytes | Panlaodb | ENSG00000254087.7 | 0.514039744 | 1.84706959 | 80.51858987 | 0 | | | | |
| MS4A7 | Monocytes | Panlaodb | ENSG00000166927.12 | 0.225566709 | 1.332704168 | 61.82124523 | 0 | | | | |
| CEBPB | Monocytes | Panlaodb | ENSG00000172216.5 | 0.522856989 | 0.939328506 | 34.66610271 | 1.4441E-252 | | | | |
| IFI30 | Monocytes | Panlaodb | ENSG00000216490.3 | 0.141886429 | 1.912730045 | 90.1358298 | 0 | | | | |
| SERPINA1 | Monocytes | Panlaodb | ENSG00000197249.13 | 0.350227401 | 0.978183189 | 44.21446572 | 0 | | | | |
| RGS2 | Monocytes | Panlaodb | ENSG00000116741.7 | 0.562267114 | 1.032831901 | 39.1448344 | 6.159823e-318 | | | | |
| LST1 | Monocytes | Panlaodb | ENSG00000204482.10 | 0.225127943 | 1.472930535 | 63.1668714 | 0 | | | | |
| SPI1 | Monocytes | Panlaodb | ENSG00000066336.11 | 0.374760974 | 0.758601477 | 28.10151546 | 5.0997E-169 | | | | |
| TYMP | Monocytes | Panlaodb | ENSG00000025708.13 | 0.233216449 | 2.258428811 | 119.5950663 | 0 | | | | |
| CSTA | Monocytes | Panlaodb | ENSG00000121552.3 | 0.330812428 | 2.237390229 | 122.6893473 | 0 | | | | |
| FGL2 | Monocytes | Panlaodb | ENSG00000127951.6 | 0.384238554 | 1.280512343 | 50.5638507 | 0 | | | | |
| PYCARD | Monocytes | Panlaodb | ENSG00000103490.13 | 0.259190321 | 0.824268064 | 33.79448544 | 1.2998E-240 | | | | |
| LYST | Monocytes | Panlaodb | ENSG00000143669.13 | 0.458892682 | 1.913094513 | 87.03807778 | 0 | | | | |
| IL1B | Monocytes | Panlaodb | ENSG00000125538.11 | 0.286291138 | 1.595274938 | 66.02576165 | 0 | | | | |
| CFP | Monocytes | Panlaodb | ENSG00000126759.13 | 0.239420509 | 2.379180795 | 128.4749668 | 0 | | | | |
| CD36 | Monocytes | Panlaodb | ENSG00000135218.18 | 0.276036411 | -0.037627285 | -1.322528821 | 0.186014016 | | | | |
| CD274 | Monocytes | CellMarker | ENSG00000120217.13 | 0.077176926 | 0.08286784 | 2.905751647 | 0.003669538 | | | | |
| F3 | Monocytes | CellMarker | ENSG00000117525.13 | 0.026642647 | 1.404623209 | 63.89735759 | 0 | | | | |
| PDCD1 | Monocytes | CellMarker | ENSG00000188389.10 | | 1.374012265 | 78.616869 | 0 | | | | |
| TNFRSF1B | Monocytes | CellMarker | ENSG00000028137.18 | 0.482560131 | 0.198782588 | 7.39072047 | 1.54419E-13 | | | | |
| ITGB2 | Monocytes | CellMarker | ENSG00000160255.17 | 0.678468611 | 1.374012265 | 78.616869 | 0 | | | | |
| SECISBP2L | Monocytes | CellMarker | ENSG00000138593.8 | 0.330091472 | 0.198782588 | 7.39072047 | 1.54419E-13 | | | | |
| CXCL8 | Granulocytes | CellMarker | ENSG00000169429.10 | 0.393517967 | 1.583342584 | 94.51982742 | 0 | | | | |
| CCR3 | Granulocytes | CellMarker | ENSG00000183625.14 | 0.133266979 | 0.454935333 | 19.9220324 | 4.36208E-87 | | | | |
| ENPP3 | Granulocytes | CellMarker | ENSG00000154269.14 | 0.060834126 | -0.015063446 | -0.645801376 | 0.518418726 | | | | |
| ITGA4 | Granulocytes | CellMarker | ENSG00000115232.13 | 0.511867052 | -0.537980245 | -27.48720269 | 5.4852E-162 | | | | |
| CD63 | Granulocytes | CellMarker | ENSG00000135404.11 | 0.494453346 | -0.059986417 | -2.950808677 | 0.003174754 | | | | |
| MPO | Granulocytes | CellMarker | ENSG00000005381.7 | 0.062526423 | -0.049959033 | -2.142394375 | 0.03217919 | | | | |
| LCN2 | Granulocytes | CellMarker | ENSG00000148346.11 | 0.038019 | -0.002328798 | -0.09972695 | 0.920562552 | | | | |
| FPR1 | Granulocytes | CellMarker | ENSG00000171051.8 | 0.448202039 | 1.286503927 | 72.21967549 | 0 | | | | |
| ANPEP | Granulocytes | CellMarker | ENSG00000166825.13 | 0.269800583 | 0.444058914 | 20.01278936 | 7.46183E-88 | | | | |
| C5AR1 | Granulocytes | CellMarker | ENSG00000197405.7 | 0.473750206 | 1.275206495 | 72.82098054 | 0 | | | | |
| CXCR1 | Granulocytes | CellMarker | ENSG00000163464.7 | 0.305855534 | 1.770712053 | 108.0922096 | 0 | | | | |
| CXCR2 | Granulocytes | CellMarker | ENSG00000180871.7 | 0.379957926 | 2.157967033 | 188.7986038 | 0 | | | | |
| JAML | Granulocytes | CellMarker | ENSG00000160593.18 | 0.400921826 | 0.593366523 | 28.50092892 | 1.1527E-173 | | | | |
| TLR2 | Granulocytes | CellMarker | ENSG00000137462.7 | 0.251485333 | 0.508171217 | 22.88461144 | 8.5399E-114 | | | | |

**Description: the words marked by red color in this table indicate that the corresponding gene is significantly differentially expressed in bulk-seq data or scRNA-seq data; The differential expressed gene analysis was conducted by limma.**
